# Determinants of Motor Neuron Functional Subtypes Important for Locomotor Speed

**DOI:** 10.1101/2022.12.23.521820

**Authors:** Kristen P. D’Elia, Hanna Hameedy, Dena Goldblatt, Paul Frazel, Mercer Kriese, Yunlu Zhu, Kyla R. Hamling, Koichi Kawakami, Shane A. Liddelow, David Schoppik, Jeremy S. Dasen

**Affiliations:** Department of Neuroscience & Physiology, and the Neuroscience Institute, New York University Grossman School of Medicine; Department of Otolaryngology, New York University Grossman School of Medicine; Center for Neural Science, New York University; Division of Molecular and Developmental Biology, National Institute of Genetics, Mishima, Shizuoka, Japan

## Abstract

Locomotion requires precise control of the strength and speed of muscle contraction and is achieved by recruiting functionally-distinct subtypes of motor neurons (MNs). MNs are essential to movement and differentially susceptible in disease, but little is known about how MNs acquire functional subtype-specific features during development. Using single-cell RNA profiling in embryonic and larval zebrafish, we identify novel and conserved molecular signatures for MN functional subtypes, and identify genes expressed in both early post-mitotic and mature MNs. Assessing MN development in genetic mutants, we define a molecular program essential for MN functional subtype specification. Two evolutionarily-conserved transcription factors, Prdm16 and Mecom, are both functional subtype-specific determinants integral for fast MN development. Loss of *prdm16* or *mecom* causes fast MNs to develop transcriptional profiles and innervation similar to slow MNs. These results reveal the molecular diversity of vertebrate axial MNs and demonstrate that functional subtypes are specified through intrinsic transcriptional codes.

## INTRODUCTION

To modulate locomotor speed, animals engage functionally specialized motor neurons (MNs) to contract distinct muscle fibers ^1, 2^. Smaller muscle fibers, called slow-twitch, produce slower and sustained movements, while larger fast-twitch fibers are recruited progressively to increase speed or force ^3, 4^. Mammalian MNs innervate either fast or slow-twitch muscle fibers. These MN “functional subtypes” have diverse attributes matched to their roles in muscle contraction ^5^. Notably, in diseases such as amyotrophic lateral sclerosis (ALS), fast MNs are more susceptible to degeneration ^6^. Although MN functional subtypes are essential for locomotor speed control, little is known about the molecular determinants that specify their identity.

Spinal MNs can also be differentiated anatomically as they cluster into motor pools that innervate individual muscles (e.g. biceps or triceps) ^7^. In contrast to functional subtypes, the mechanisms that specify MN anatomical subtypes are well understood ^3, 7^. During embryonic development, transcription factors such as Lim-, Hox-, and Mnx-homeodomain proteins drive cascades of gene expression that determine muscle-specific targeting, somatotopic organization, and integration into pre-motor locomotor networks ^8–12^. Mammalian motor pools contain both fast and slow functional subtypes, with no clear anatomic organization, hindering an understanding of how they are specified during development. Recent transcriptional profiling of mature MNs has uncovered differentially expressed genes that mark fast and slow MNs ^13–15^, underscoring a key question: do transcriptional codes specify functional subtypes of MNs?

The larval zebrafish is an ideal model system to define the molecular determinants of MN functional subtypes. First, the molecular determinants of both appendicular (fin) and axial (trunk) motor neurons are highly conserved ^16–22^ and regulate muscle targeting ^23–25^. Second, MN functional subtype identities are mapped in both space (dorso-ventral soma position) and time (birthdate). Larval zebrafish have three broad classes of axial MNs: primary (fast, dorsal, first-born), fast-secondary (fast, dorsal, middle-born), and slow (ventral, last-born) (Figure 1A). These three classes can be further subdivided along characteristic anatomical and physiological lines ^26–33^. Finally, the exceptional genetic and optical accessibility of zebrafish facilitates *in vivo* evaluation of functional subtypes after loss-of-function, the gold standard for identifying molecular determinants.

**Figure 1:**
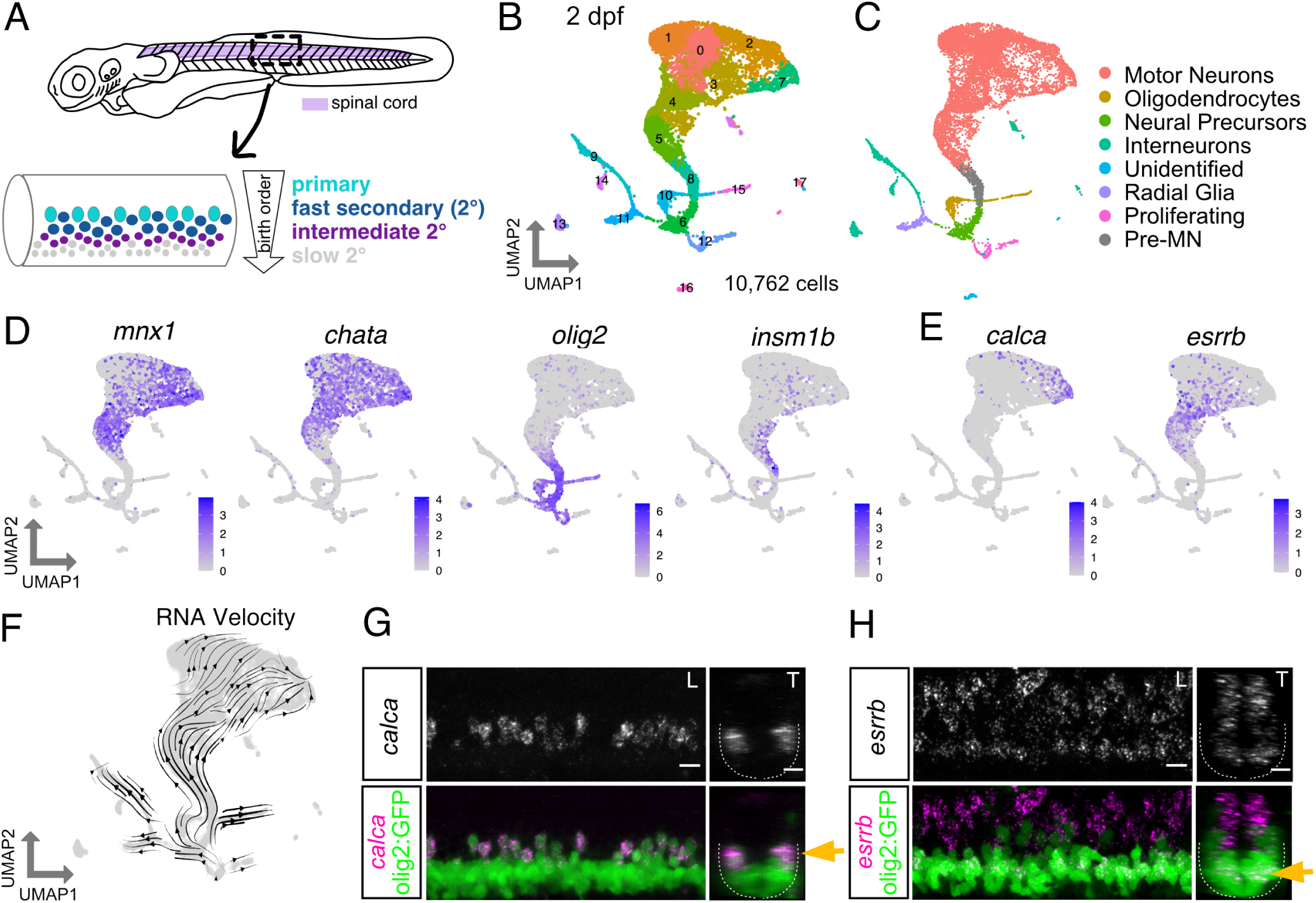
Zebrafish motor neuron functional subtypes are molecularly distinct. (A) Diagram of 3 dpf zebrafish showing location of spinal cord in purple. Crop of the tail shows the spinal cord at a lateral view. MN functional subtypes indicated by colors are organized along the dorsoventral axis and by birthdate. (B) UMAP representation of 10,762 GFP^+^ calls at 2 dpf from 90 *Tg(olig2::GFP)* larvae. Each dot represents one cell; 18 cell clusters are represented by coloring. (C) Major cell classes indicated by colors represented in the UMAP. (D) Feature plots of marker genes for MN maturity. Purple indicates expression. (E) Feature plots for markers of mammalian fast (*calca*) and slow (*esrrb*) MNs. (F) RNA Velocity streamlines overlaid on UMAP coordinates in (B). (G) FISH of *calca* at 2 dpf. Left image (L) is lateral view and right image (T) is transverse view of the spinal cord. Gold arrow indicates dorsoventral localization of gene expression within MN population. Dotted line indicates ventral spinal cord boundary. (H) FISH of *esrrb* at 2 dpf. Scale bars for G/H, 10 µm.

Here we leverage the organization of MNs in zebrafish to (1) uncover the molecular diversity among MN functional subtypes and (2) identify transcriptional determinants of MN functional subtype identity. First, we discovered remarkable molecular diversity among fast and slow secondary MNs, and defined both conserved and novel markers that are maintained between early developmental stages through functional maturity. Next, we focused on two conserved transcription factors expressed in fast secondary MNs: Prdm16 and Mecom. Loss of these genes leads fast secondary MNs to acquire the anatomical projections and molecular features that define slow MNs. Our data provide novel insights into the molecular diversity of vertebrate spinal MNs, and establish that MN functional subtypes are determined by intrinsic genetic codes acting early in development.

## RESULTS

### Molecular diversity of zebrafish motor neuron subtypes

We first set out to evaluate the molecular heterogeneity of zebrafish MNs during development at 2 days post fertilization (dpf) (Figure S1A). At 2 dpf, the oldest MNs (primary MNs and fast secondary MNs) are largely mature and active during locomotion ^34–37^. At this time point, slow secondary MNs have been recently born and are in the process of innervating muscle. To isolate MNs we used a *Tg(olig2:DsRed2)^vu19^* reporter line that labels the pMN progenitor domain and its progeny (which includes MNs, oligodendrocytes, and some interneurons) ^38, 39^. After dissection of the body caudal to the hindbrain, dissociation, and isolation of cells by fluorescence-activated cell sorting (FACS), we performed single cell RNA sequencing (scRNAseq) (Figures S1B and S1H). We characterized a total of 10,762 cells after filtering (Figures 1 and S1). Analyses using known markers identified several broad classes of cell types among clusters (Figures 1C and 1D) including MNs, oligodendrocytes, and interneurons (Figures S1C and S1D). MN clusters were identified as those expressing *chata* (Figure 1D).

As birthdate and functional subtype are linked in zebrafish MNs, we determined the relative birth order of different MN clusters and their respective putative functional class (Figure 1A). The expression of two genes, *olig2* and *insmb1*, which are progenitor and pro-neural genes respectively, suggested clusters 5 and 8 were younger MNs (Figures 1B and 1D). The expression of mature MN marker (*chata*) was higher in MN clusters 0, 1, 2, 3, 4, and 7 indicating they are likely more mature MNs. We used RNA Velocity ^40, 41^ to assay the progression of gene expression profiles across development. RNA Velocity streamlines diverged from the progenitor population into clusters that include four major cell types (MNs, oligodendrocytes, interneurons, and radial glia) (Figures 1C and 1F). We integrated our data with a previously published dataset of *olig2*^+^ MN populations at 1 dpf and 1.5 dpf ^39^. We found these datasets could be well integrated (Figure S1E). RNA Velocity on the merged datasets appropriately predicted trajectories and pseudotime based on sample age (Figures S1F and S1G). With well-defined MN subtype birthdates, cluster birth order determined by RNA Velocity implies we can potentially differentiate young (slow) from old (fast) functional subtypes.

We investigated whether previously characterized mammalian markers of adult fast and slow MNs could discriminate functional subtypes in embryonic zebrafish MNs. We observed expression of a fast MN marker, *calca*, was restricted to putative older MN clusters while expression of a slow MN marker, estrogen-related receptor beta (*esrrb* in zebrafish), was enriched in putative younger MN clusters (Figure 1E) ^42^. This segregation is consistent with these markers being conserved in fast and slow zebrafish MN functional subtypes, respectively. We acknowledge the limits of RNA Velocity ^43^, and therefore confirmed the relative birthdates of these clusters by determining the localization of these putative MN functional subtype markers *in vivo* by fluorescent *in situ* hybridization (FISH). This analysis showed dorsal MNs express *calca* and ventral MNs express *esrrb* (Figures 1G, 1H, S5A and S5B). These results also demonstrated the inferred ages of MN clusters from the RNA Velocity and maturity analysis were consistent with marker localization.

To further investigate the molecular diversity among secondary MNs, we performed a deeper analysis of *chata*^+^ MN clusters at 2 dpf (6,103 MNs; Figure 2A). A heatmap of marker genes showed clear transcriptional heterogeneity across *chata*^+^ clusters (Figure S2E). Genes which distinguished clusters included transcription factors, guidance molecules, signaling and structural proteins. A Gene Ontology analysis of MN cluster markers revealed a particular enrichment for genes involved in neural development and metabolism (Figure S2F). To determine if the detected molecular diversity at 2 dpf was only due to the birthdate differences of the subtypes or if it reflected sustained genetic differences, we assessed MN molecular diversity at 5 dpf when all functional subtypes have innervated and can contract muscle ^27^. We performed the analogous experiment (Figure S2B) at 5 dpf assessing 14,882 cells after filtering (Figure S2A). We similarly repeated our analyses on *chata*^+^ MN clusters at 5 dpf (8,087 MNs; Figures S2A to S2C and S2E). Comparing the two datasets (Figures 2A and 2B), 74% of cluster marker genes in the 5 dpf data set were shared with the 2 dpf dataset, indicating the majority were conserved across timepoints (Figure 2C). We also performed a correlation analysis between the clusters at the two timepoints. Assessing intergroup distances among clusters at both 2 dpf and 5 dpf, we found most of the clusters at the earlier stage were closely related to a corresponding 5 dpf cluster (Figure 2D). These results indicate the molecular diversity between subtypes at 2 dpf reflects sustained molecular differences among subtypes.

**Figure 2:**
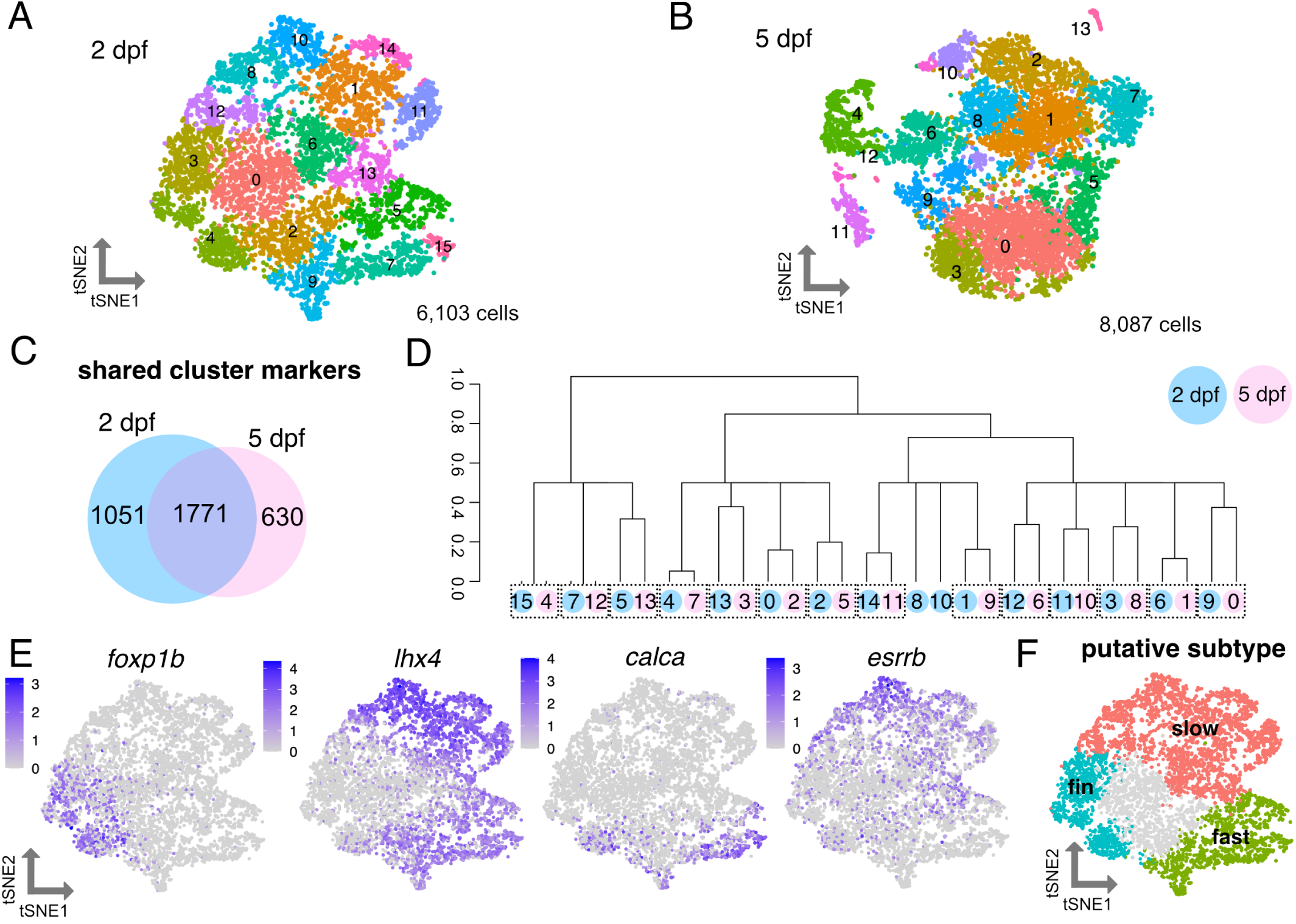
Heterogeneity of motor neuron functional subtypes is identifiable during development and at functional maturity. (A) tSNE representing 6,103 MNs at 2 dpf. (B) tSNE representing 8,087 MNs at 5 dpf. (C) Venn diagram demonstrating overlap in cluster markers from 2 dpf and 5 dpf MNs. (D) Dendrogram demonstrating relationships between 2 dpf and 5 dpf MN clusters. Circle color indicates age (blue for 2 dpf, pink for 5 dpf). Circle number indicates cluster from A or B. Dotted boxes group a 2 dpf cluster and its most closely related 5 dpf cluster. Expression of *Hox* genes may also contribute conserved molecular profiles between the 2 dpf and 5 dpf datasets. (E) Feature plots showing fin (*foxp1b*), axial (*lhx4*), fast (*calca*), and slow (*esrrb*) MN marker expression within 2 dpf MN clusters. Purple indicates expression. (F) General map within tSNE space of putative MN subpopulations within 2 dpf MN clusters.

### Conserved and divergent expression of motor neuron subtype markers

To delineate which MN subtype each cluster represents, we first evaluated gene expression of known determinants of subtype fates. We found previously that a marker of limb-innervating MNs, *Foxp1*, is expressed by pectoral fin MNs in zebrafish larvae at 3 dpf and these MNs lacked expression of an axial MN determinant, *lhx3* ^44^. We marked clusters within the 2 dpf data (Figure 2E) that expressed *foxp1b* and lacked expression of *lhx3* and *lhx4* as pectoral fin MNs (Figure S2G). Expression of an adductor fin MN subtype marker, *lhx1a* ^45^, marked a portion of these clusters (Figure S2G). *lhx3/4* were widely expressed within almost all remaining axial MN clusters, in contrast with tetrapod axial MNs where expression of *Lhx3* is sustained only by dorsally-projecting medial motor column (MMC) axial MNs.

Axial MN clusters could be classified as putative fast or slow axial MNs based on expression of *calca* and *esrrb* (Figures 2E and 2F). These broad functional groups include multiple clusters, indicating molecular diversity that could reflect additional subtype divisions. For example, MNs within the fast and slow groups could be distinguished by differences in *Hox* gene expression, likely reflecting the rostrocaudal domain from which they originated (Figure S2H). Based on these marker assessments, we assigned putative broad functional identities of the clusters (Figure 2F).

### Molecular signatures of zebrafish motor neuron functional subtypes

In addition to *calca* and *essrb*, we identified and characterized several novel markers of axial MN subtypes (Figures S3 and 3A). We confirmed the localization of expression of novel markers using FISH in the *Et(-0.6hsp70l:GAL4-VP16)^s^*^1020t^;*Tg(UAS:GFP)* line, hereafter referred to as *Tg(olig2::GFP)*, which labels the pMN domain and its progeny including all MNs. We determined general dorsal-ventral soma positioning of mature MNs using Mnx (Hb9) antibody staining (Figure 3B). The most ventral GFP^+^ cells were Mnx^-^ indicating they were progenitors or oligodendrocytes. Within the mature population, dorsal MNs were generally classified as fast and the ventral MNs were generally classified as slow. We did not classify intermediate MNs since their subtype specificity has not been determined at larval stages. We found multiple novel markers expressed in fast MNs including *pcp4a*, *calb2b*, *nefma*, *prdm16*, *scn4ba*, *gfra1b*, *mafba*, and *rbfox3a* (Figures 3A and S5). *pcp4a* and *calb2b* labeled the dorsal-most fast axial MNs including both primary and secondary MNs (Figures 3C and 3D). These markers also maintained their dorsal expression at 5 dpf (Figures S4A and S4B).

**Figure 3:**
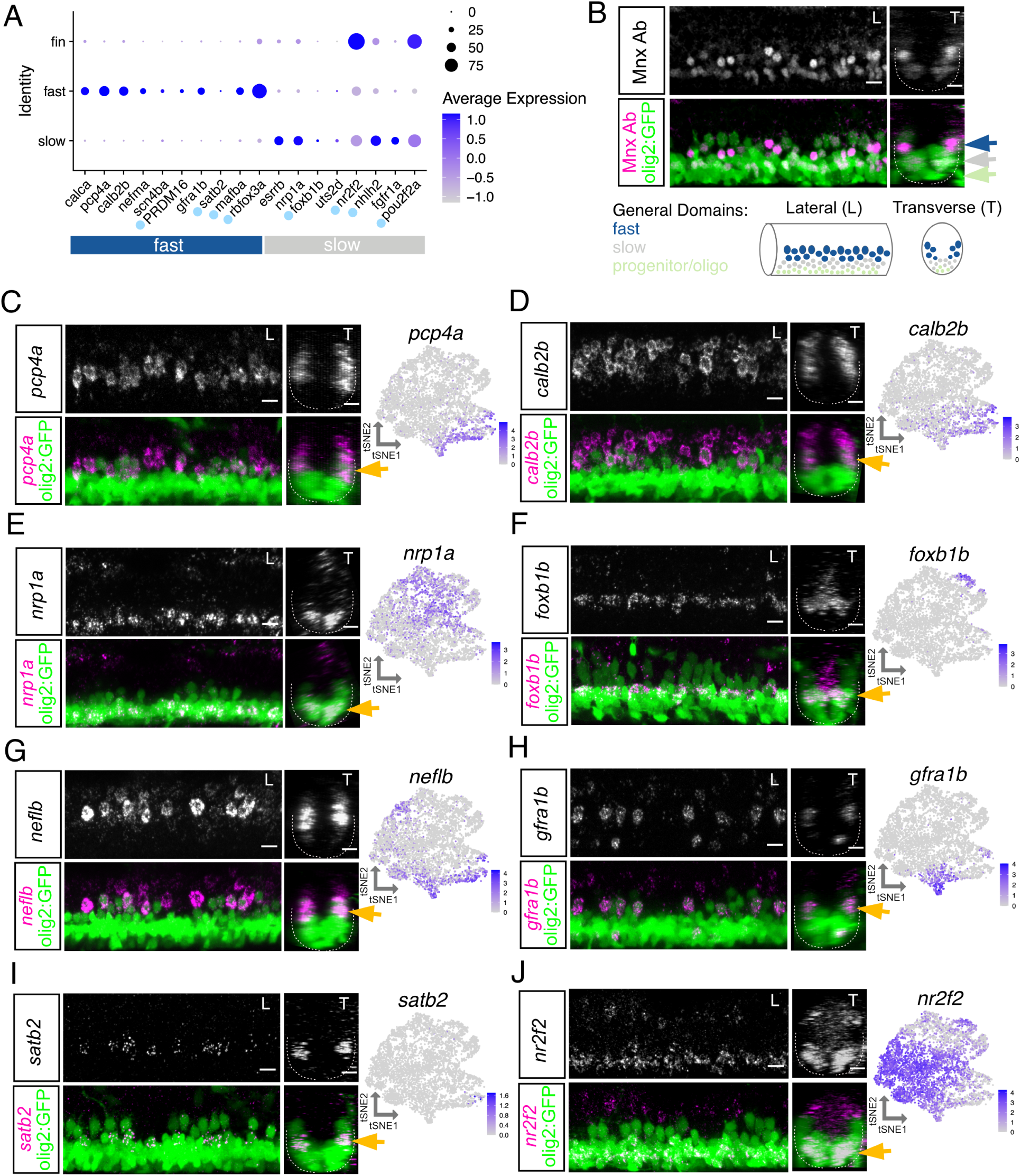
Characterization of motor neuron functional subtype-restricted markers. (A) Dot plot indicating expression of general fast and slow MN functional subtype markers. Light blue dot next to gene name indicates a transcription factor. At 2 dpf uts2d is expressed by a small number of MNs. (B) Top: Antibody stain demonstrating domain of Mnx (Hb9) expression within mature MNs in *Tg(olig2::GFP)* larvae at 2 dpf. Bottom: Schematic indicating general domains of expression considered when verifying functional subtype expression. Dorsal GFP+ cells are putative fast MNs (blue), most ventral are progenitors or oligodendrocytes (green), and those between are putative slow MNs (gray). (C-J) FISH and feature plots for MN subpopulation markers at 2 dpf. Gold arrow indicates dorsoventral localization of FISH expression within MN population. Dotted line indicates ventral spinal cord boundary. Left image (L) is lateral view and right image (T) is transverse view. Scale bars, 10 µm.

While many markers of slow MN clusters were shared with fin MNs (*nr2f2*, *nhlh2*, and *pou2f2a*), we found several that had specific slow MN expression including *nrp1a*, *foxb1b*, *uts2d*, and *fgfr1a* (Figure 3A). In addition to transcription factors, cluster-restricted genes encode molecules that are related to axon guidance, growth, and signaling. *nrp1a*, which has a role in axon guidance ^46^, was widely expressed within all slow clusters and, accordingly, expressed within ventral MNs (Figures 3E and S5). Some markers, such as *uts2d*, were reported to be present in MNs ^47^, but their subpopulation-specificity and function remain unknown. We did not identify any primary MN-specific markers within our dataset, but found many of the fast MN markers labeled both primary and secondary dorsal MNs. Some of these markers had higher expression in primary MNs such as *neflb* (Figures 3G and S5).

Several novel markers labeled smaller subsets of functional groups including *gfra1b* which labeled two of the fast MN clusters and *foxb1b* which labeled two slow MN clusters (Figures 3F and 3H). FISH at 2 dpf revealed *gfra1b* was expressed in only a subset of dorsal fast MNs. We found an array of guidance molecules with cluster-specific expression that could potentially contribute to between- and within-subtype innervation diversity (Figures S4C and S4D). The restricted expression of *foxb1b* within two slow MN clusters was reflected in its more limited expression within the ventral domain. *foxb1b* did not label as many of the most ventral MNs as did the slow MN marker *nrp1a*; *foxb1b* might therefore label MNs involved in intermediate speed swimming.

We also identified markers of mammalian axial MN subpopulations that are conserved in zebrafish. A recent study ^48^ found subpopulations of tetrapod axial MMC neurons marked by *Satb2*, *Nr2f2*, and *Bcl11b*. We found all three markers expressed within zebrafish axial MNs. *satb2* was restricted within one cluster and confirmed to be in only a few MNs per hemisegment by FISH (Figure 3I). *nr2f2* was strongly expressed within fin MN clusters and a portion of axial MNs. FISH demonstrated *nr2f2* to be enriched in ventral axial MNs (Figure 3J). *bcl11ba* was detected in both fast and slow axial MN clusters and labeled all spinal cord cell types by FISH (data not shown). This data demonstrates that tetrapod axial MN subtype markers are conserved in zebrafish, though their subtype restrictions likely vary.

### Prdm16 and Mecom mark putative fast axial motor neurons

To identify potential fate-determining genes for MN functional subtypes, we assessed transcription factors that were expressed in either fast or slow clusters (Figure 3A). A transcription factor from the Prdm family, *prdm16*, was detected in fast MN clusters and is a paralog of a previously described axial MMC-specific transcription factor in mice, *Mecom* (also known as *Evi1* or *Prdm3*) ^49^. Both *prdm16* and *mecom* were expressed in fast MN clusters at 2 dpf (Figure 4A). *In situ* hybridization at 30 hpf revealed that both *prdm16* and *mecom* are expressed in a clustered population of cells in the ventral spinal cord (Figure 4B). Using an antibody that recognizes both Prdm16 and Mecom (Figures 4C and 4D magenta), as well as one that recognized Mecom alone (Figure 4D green), we confirmed these transcription factors were expressed in a subset of dorsal MNs at 2 dpf. We also detected *prdm16* transcripts in mouse MMC neurons (Figure S6A), as well as Mecom protein in chick and skate axial MNs (Figures S6B and S6C), indicating expression of these factors is evolutionarily conserved.

**Figure 4:**
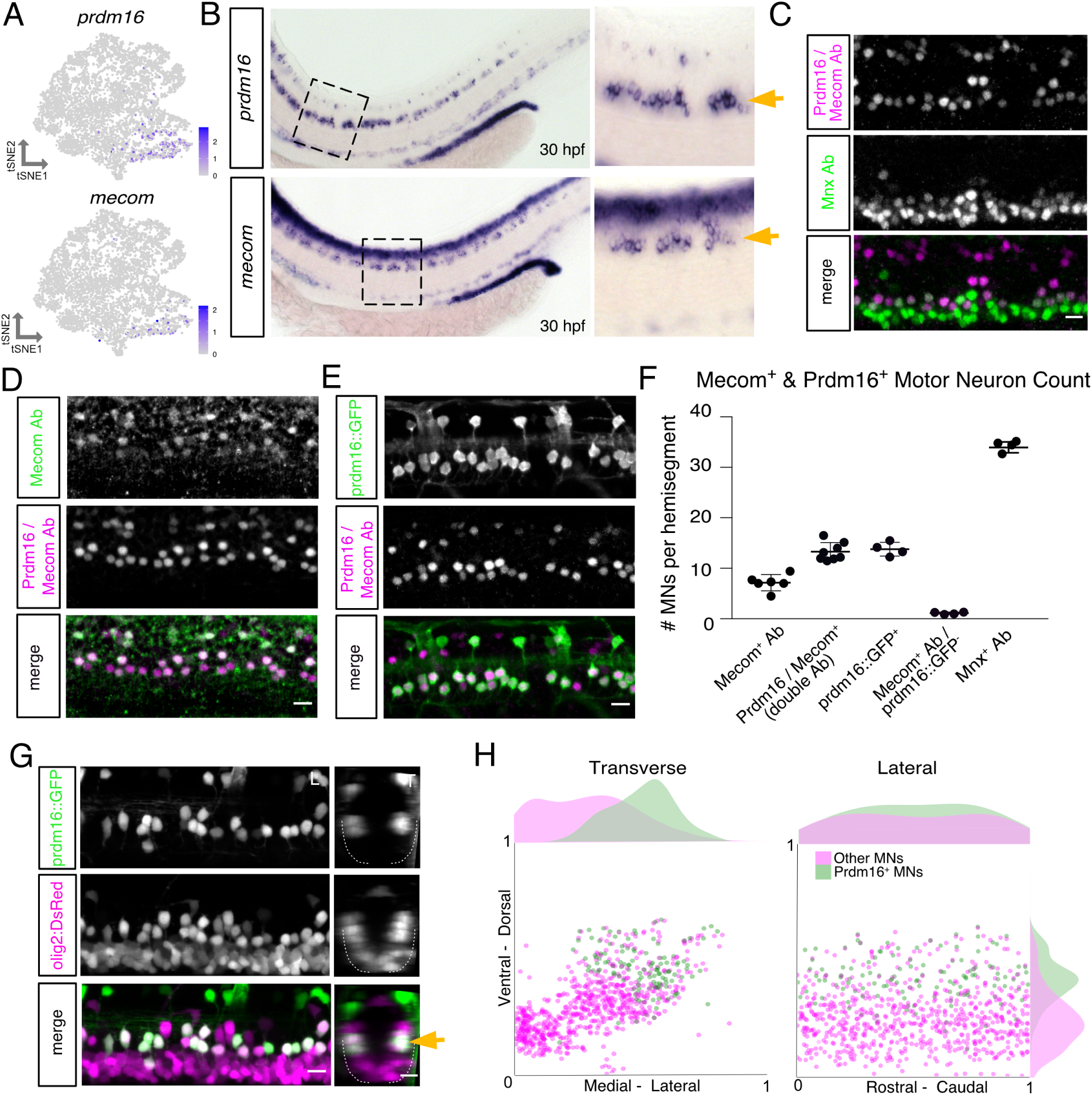
Prdm16 and Mecom mark dorsally-positioned motor neurons. (A) Feature plot of *prdm16* and *mecom* expression in 2 dpf MN tSNE showing localization in fast MN clusters as in Figure 2F. (B) *prdm16* and *mecom in situ* hybridization at 30 hpf in zebrafish. Zoomed image indicated by dashed box. Gold arrows indicate MNs. Additional staining in the dorsal spinal cord is labeling of some interneurons (*prdm16*) and a progenitor domain (*mecom*). (C) Antibody stain of Prdm16/Mecom within Mnx^+^ MNs in the zebrafish spinal cord at 2 dpf. Dorsal Prdm16/Mecom^+^; Mnx^-^ cells are a subpopulation of interneurons. (D) Antibody stain showing overlap in expression between Prdm16/Mecom double antibody and Mecom alone antibody at 2 dpf. (E) Antibody stain showing overlap in expression between GFP^+^ cells in *Tg(prdm16::GFP)* line and Prdm16/Mecom double antibody at 2 dpf. (F) Quantification of Prdm16 and/or Mecom-expressing MNs from stains in (C-E). Line with bars indicates average with SEM. (G) Image of the spinal cord of a *Tg(prdm16::GFP)*;*Tg(olig2:DsRed2)* larvae at 5 dpf. Gold arrow indicates dorsoventral localization of GFP^+^ MNs within DsRed2^+^ population. Left image (L) is lateral view and right image (T) is transverse view. (H) Soma position plot for GFP^+^ MNs and all other MNs quantified from five larvae at 5 dpf as in (G). Left plot shows the transverse view. Right plot shows the lateral view. Histograms on the top and sides of the plots indicate the distribution of cells across the axes. Green indicates GFP^+^;DsRed2^+^ cells. Magenta indicates GFP^-^;DsRed^+^ cells. While magenta (Other MNs) count could include *olig2* progenitors or oligodendrocytes, these were largely excluded based on morphology and localization. Scale bars, 10 µm.

To visualize Prdm16^+^ MNs, we crossed a GAL4 enhancer trap at the *prdm16* locus *gSAIzGFFM1116A*, hereafter referred to as *Tg(prdm16:GAL4)*, with a *Tg(5xUAS:EGFP)* line, hereafter called *Tg(prdm16::GFP)*. Staining 2 dpf embryos with the Prdm16/Mecom double antibody demonstrated near complete overlap with GFP^+^ MNs (∼92%; Figure 4E). Cell counting indicated that there are ∼14 Prdm16^+^ positive cells per hemisegment, and about half were also Mecom^+^ (Figure 4F). We crossed the *Tg(prdm16::GFP)* line to an *Tg(olig2:DsRed2)* line, and found the dorsal positioning of GFP^+^ MNs was maintained until 5 dpf (Figure 4G). There was a reduction of about 3 or 4 GFP^+^ MNs per hemisegment from 3 dpf to 5 dpf, possibly reflecting a decay in fluorescence in the oldest MNs (Figure S7A). By normalizing the position of Prdm16^+^ MN soma to landmarks (see Methods), we observed they make up most, but not all, dorsal and lateral MNs at 5 dpf (Figure 4H). The remaining dorsal GFP^-^ MNs are likely primary MNs due to their large size and positioning. Consistent with the scRNAseq data, expression patterns of Prdm16 and Mecom suggest they mark secondary MNs involved in fast swimming.

### Prdm16 marks secondary motor neurons that innervate fast muscle

In zebrafish, slow muscle is located superficially and innervated by septal (s) nerves, while deeper fast muscle is innervated by medial (m) nerves (Figure 5A) ^31^. To determine which muscle fiber types are innervated by Prdm16^+^ MNs, we performed confocal microscopy on *Tg(prdm16::GFP)*;*Tg(olig2:DsRed2)* fish. The entire MN population, represented by DsRed+ axons, innervated both fast and slow muscle (Figure 5B). By contrast, GFP^+^ axons primarily innervated fast muscle via the medial nerves, and the majority of DsRed^+^ axons in slow muscle were GFP^-^. A detailed quantification of innervation demonstrated Prdm16^+^ MNs innervated all four quadrants of fast muscle with a slight ventral bias (Figure 5C). These data demonstrate that Prdm16^+^ MNs predominately innervate medial fast muscles. Notably, unlike in mammals, the Prdm16^+^ MNs in zebrafish do not exclusively innervate dorsal axial musculature.

**Figure 5:**
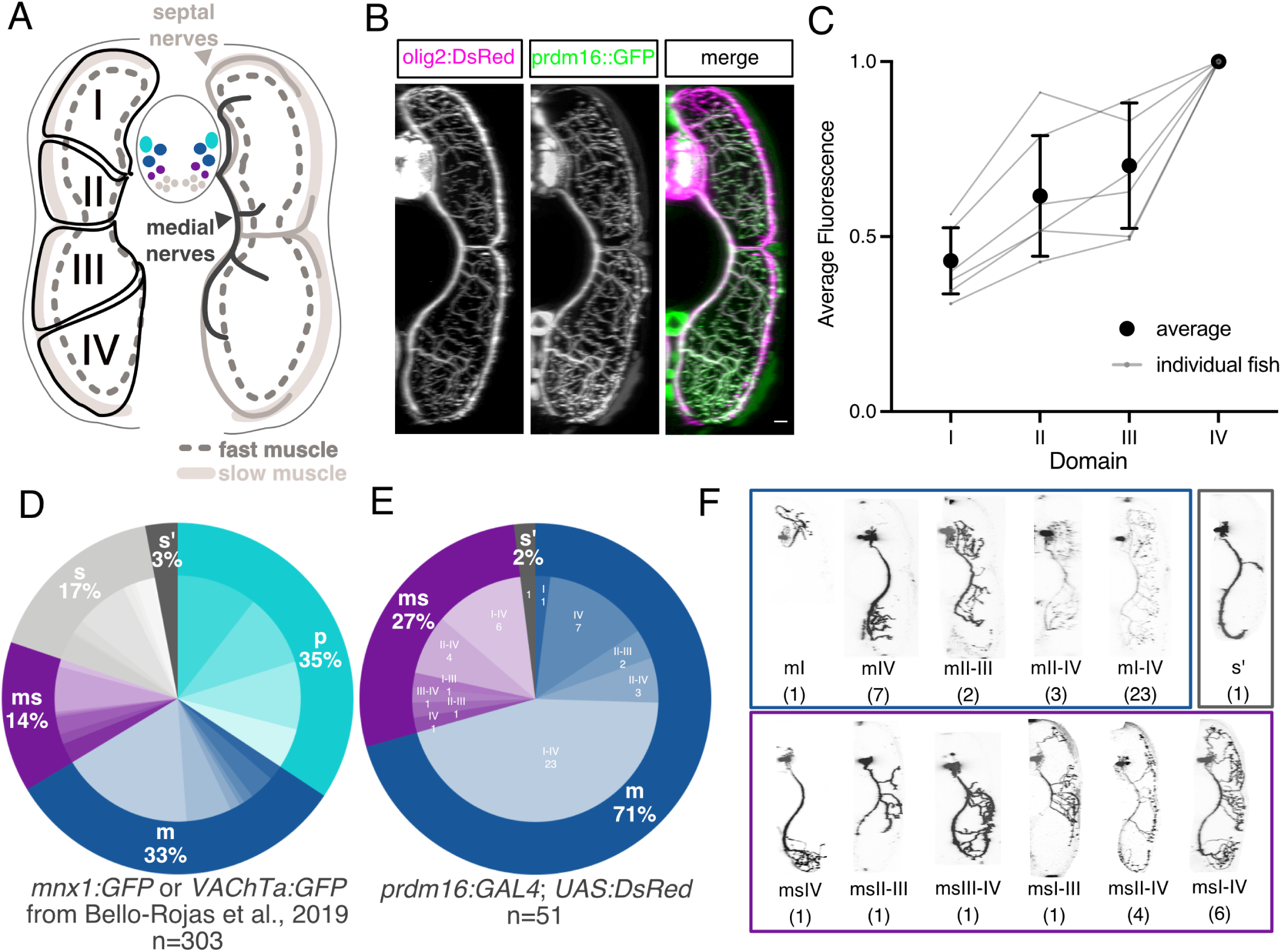
Prdm16 and Mecom define secondary fast motor neurons. (A) Schematic of a transverse cross section of the zebrafish tail showing (left) the four dorsal-ventral muscle quadrants (black solid line with labels I, II, III, IV) as defined by primary MN innervation pattern and the different muscle functional fiber types (solid light gray as slow, dashed dark gray outline as fast) and (right) the nerves that innervate them (slow muscle by septal nerves (light gray) and fast muscle by medial nerves(dark gray)). Primary MNs (p-type, light blue), responsible for escape behaviors, project along the medial nerve to innervate one of four fast muscle quadrants. m-type secondary MNs (dark blue) innervate fast muscle via the medial nerves. ms-type secondary MNs (purple) innervate both fast and slow muscle via the medial and septal nerves. s-type secondary MNs (light gray) innervate slow muscle via the septal nerves. Each individual secondary MN can innervate from one to four quadrants (I-IV) of muscle. (B) Transverse view of the zebrafish tail showing muscle innervation in *Tg(prdm16::GFP)*;*Tg(olig2:DsRed2)* larvae at 5 dpf. Scale bar, 10 µm. (C) Average GFP fluorescence from *Tg(prdm16::GFP)* within each muscle quadrant normalized to the fourth quadrant within each fish at 5 dpf. Gray lines indicate individual fish. Black points indicates average with SEM. (D) Pie chart showing subtype proportions labeled in MN sparse label experiment in ^31^. Inner subtype distribution numbers detailed in Table 1. (E) Pie chart showing subtype proportions labeled in *Tg(prdm16:GAL4)*. (F) Example images of all sparsely labeled subtypes in (E). Number in parentheses indicates number of times subtype was labeled.

**Table 1:** Sparse Label Counts. All classified MNs were *pUAS:dsRed*^+^ on a *Tg(prdm16::GFP)* background.

| Subtype | All MNs<sup>31</sup> | Prdm16+ (WT) | *mecom* -/- | *prdm16* -/- | *prdm16* -/-<br>;*mecom* +/- |
| --- | --- | --- | --- | --- | --- |
| p-type |  |  |  |  |  |
| I | 32 |  |  |  |  |
| II | 29 |  |  |  |  |
| III | 27 |  |  |  |  |
| IV | 16 |  |  |  |  |
| m-type |  |  |  |  |  |
| I | 6 | 1 |  |  |  |
| II |  |  |  |  |  |
| III |  |  |  |  |  |
| IV | 10 | 7 | 5 | 3 | 1 |
| I-II |  |  |  |  |  |
| II-III | 4 | 2 | 3 | 4 |  |
| III-IV | 2 |  |  |  |  |
| I-III | 4 |  | 3 | 1 |  |
| II-IV | 18 | 3 | 2 | 4 | 4 |
| I-IV | 53 | 23 | 12 | 14 | 5 |
| ms-type |  |  |  |  |  |
| I | 8 |  |  |  |  |
| II |  |  |  |  |  |
| III |  |  |  |  |  |
| IV | 4 | 1 | 4 |  | 2 |
| I-II |  |  |  |  |  |
| II-III | 7 | 1 | 2 |  | 2 |
| III-IV | 1 | 1 | 1 | 3 |  |
| I-III | 1 | 1 | 3 | 2 |  |
| II-IV | 18 | 4 | 4 | 6 | 2 |
| I-IV | 3 | 6 | 3 | 4 | 3 |
| s-type |  |  |  |  |  |
| I | 11 |  |  |  |  |
| II | 1 |  |  |  |  |
| III | 5 |  |  |  | 1 |
| IV | 21 |  | 1 |  | 5 |
| I-II | 1 |  |  |  |  |
| II-III | 2 |  |  |  |  |
| III-IV | 2 |  | 1 | 2 | 1 |
| I-III |  |  |  |  |  |
| II-IV | 7 |  | 4 | 7 | 3 |
| I-IV | 1 |  | 2 |  |  |
| s'-type |  |  |  |  |  |
|  | 9 | 1 | 2 |  |  |
| unclassifiable |  |  |  |  |  |
|  |  |  | 1 | 1 | 3 |
| Totals: | 303 | 51 | 53 | 51 | 32 |

**Table 2:**
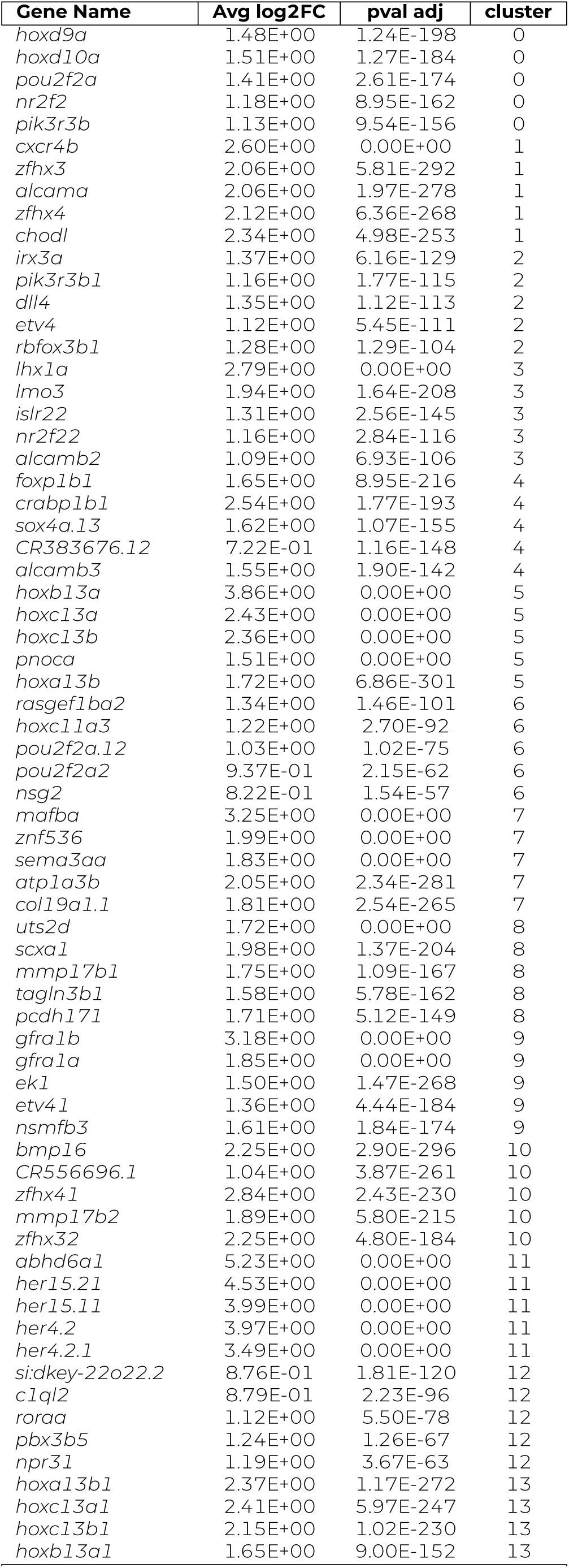
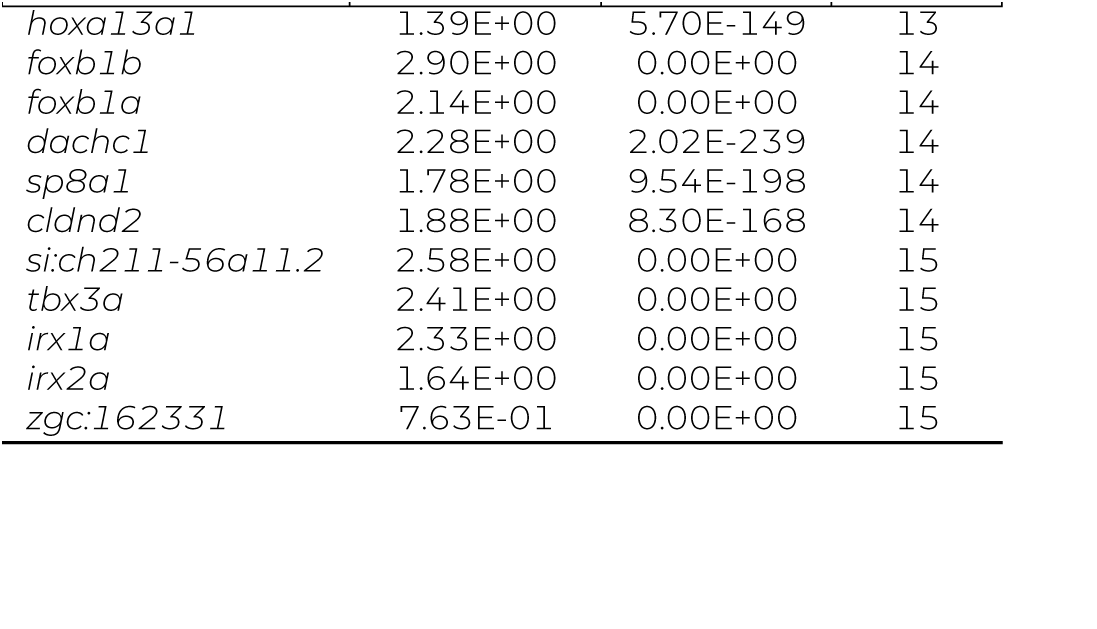
Top 5 differentially expressed genes by cluster for 2dpf MNs.

**Table 3:**
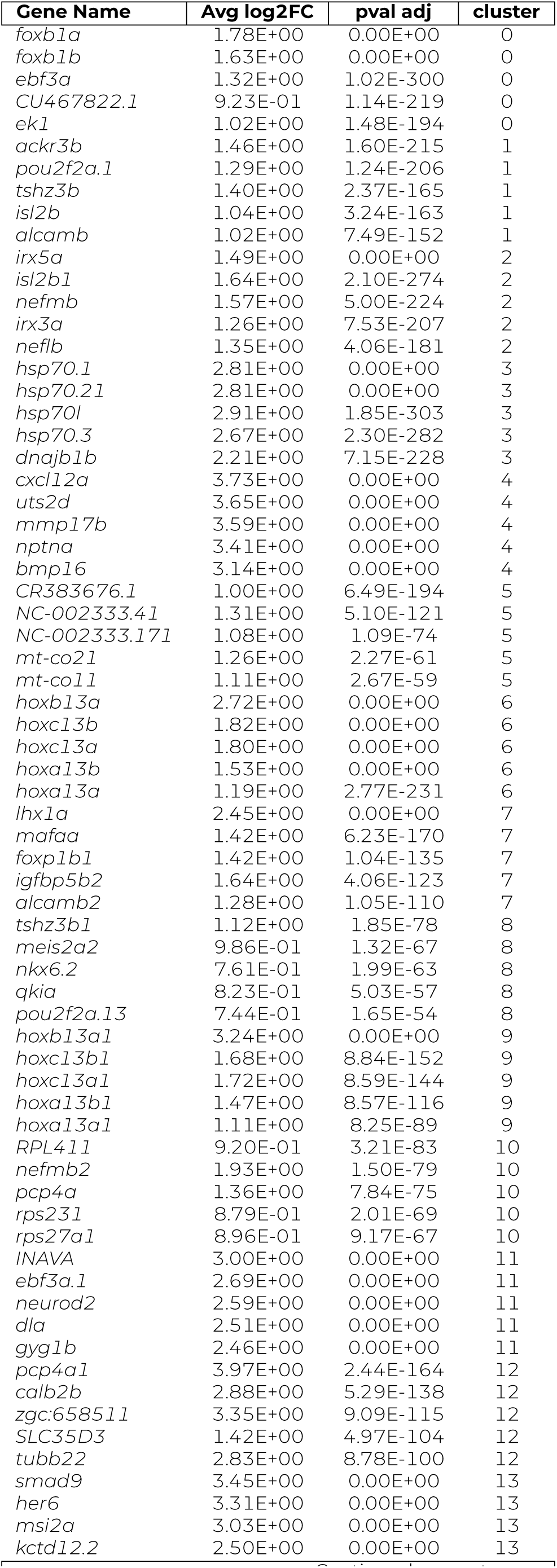
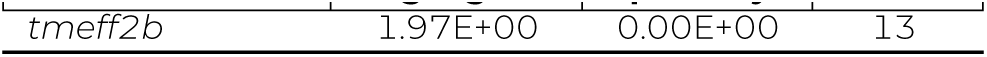
Top 5 differentially expressed genes by cluster for 5dpf MNs.

To examine the anatomical projections of single, sparsely-labeled Prdm16 MNs, we injected a *pUAS:DsRed* plasmid ^50^ into *Tg(prdm16:GAL4)* embryos at the single cell stage. Labeled cells were categorized as either fast primary MNs (p) or secondary MNs that project along the medial (m), septal (s), or both nerves (ms). Secondary MNs projecting along the medial nerve (m-type) innervate fast muscle, stype MNs innervate slow muscle, while ms-type MNs innervate both fast and slow muscle. Combining the functional class (p, m, ms, or s) and the dorsal-ventral quadrants of muscle innervation (I, II, III, and/or IV) gives a subtype classification to individual MNs in each hemi-segment (Figure 5A, Table 1). A previous study ^31^ identified 26 unique types using pan-MN labeling plasmids (Figure 5D, Table 1, n=303). We found that 98% of our labeled Prdm16^+^ MNs belonged to either the m (71%) or ms class (27%) (Figures 5E and 5F, n=51). Within these classes, there was a range of innervation subtypes (Figure 5F, Table 1). We identified 5 of 7 m-type subtypes and 5 of 7 ms-type subtypes identified previously ^31^, as well as one additional new ms subtype (ms III-IV). We labeled no p-type cells and one s’-type cell with an underdeveloped axon. A larger proportion of labeled MNs innervated the ventral quadrants, consistent with the trend detected with total population fluorescence (Figure 5C and Figure S7B). The current limits of our tools do not allow us to identify which of these Prdm16^+^ subtypes also express Mecom. Hypothesis testing by resampling (Methods) allowed us to reject the null hypothesis that Prdm16 labels all motor neurons: m (p=0.0015), p (p=0), and s (p=0) counts were all different than expected. These experiments confirm that Prdm16^+^ selectively labels m-type and ms-type fast MNs.

### Prdm16 and Mecom are essential for the specification of fast motor neurons

To determine if Prdm16 and Mecom are necessary for the specification of fast MNs in zebrafish, we adopted a loss-of-function approach. Using CRISPR/Cas9, we created *prdm16* and *mecom* mutant lines (Figures S8A to S8F). Aside from a failure to inflate their swim bladder, *prdm16* and *mecom* mutants are grossly morphologically indistinguishable from controls through 7 dpf (Figures S8C and S8D).

We first determined if loss of *prdm16* and *mecom* led to a change in MN survival or positioning. Quantification of GFP^+^ MNs in the *Tg(prdm16::GFP)*;*Tg(olig2:DsRed2)* line showed no MN loss in either *prdm16* or *mecom* mutants at 5 dpf (Figure S8G). Double mutant embryos also showed no significant change in the number of GFP^+^ MNs (Figure S8I). The position of both GFP^+^ and GFP^-^ MNs were unchanged in either *prdm16* or *mecom* mutants (Figure S8H). These results indicate *prdm16* and *mecom* are not required for fast MN survival or soma positioning.

To assess if loss of *prdm16* or *mecom* affects fast or slow muscle innervation, we analyzed the *Tg(prdm16::GFP)*; *Tg(olig2:DsRed2)* line in *prdm16* and *mecom* mutants. We observed no gross denervation of any muscle type or quadrant (Figures 6A and S9C). The innervation of each dorso-ventral quadrant shifted away from the ventral bias found in wildtype (WT) innervation, visible as a slope change in both *prdm16* and *mecom* mutants (Figure 6B). Double mutant larvae, which are overall smaller than their control and single mutant siblings, qualitatively had reduced muscle size and a decrease in fast muscle innervation (Figure S8J). These results suggest that, while fast muscle is still innervated after deletion of *prdm16* or *mecom*, there appears to be a shift in the proportion of MN subtypes and more dramatic losses with loss of both alleles.

**Figure 6:**
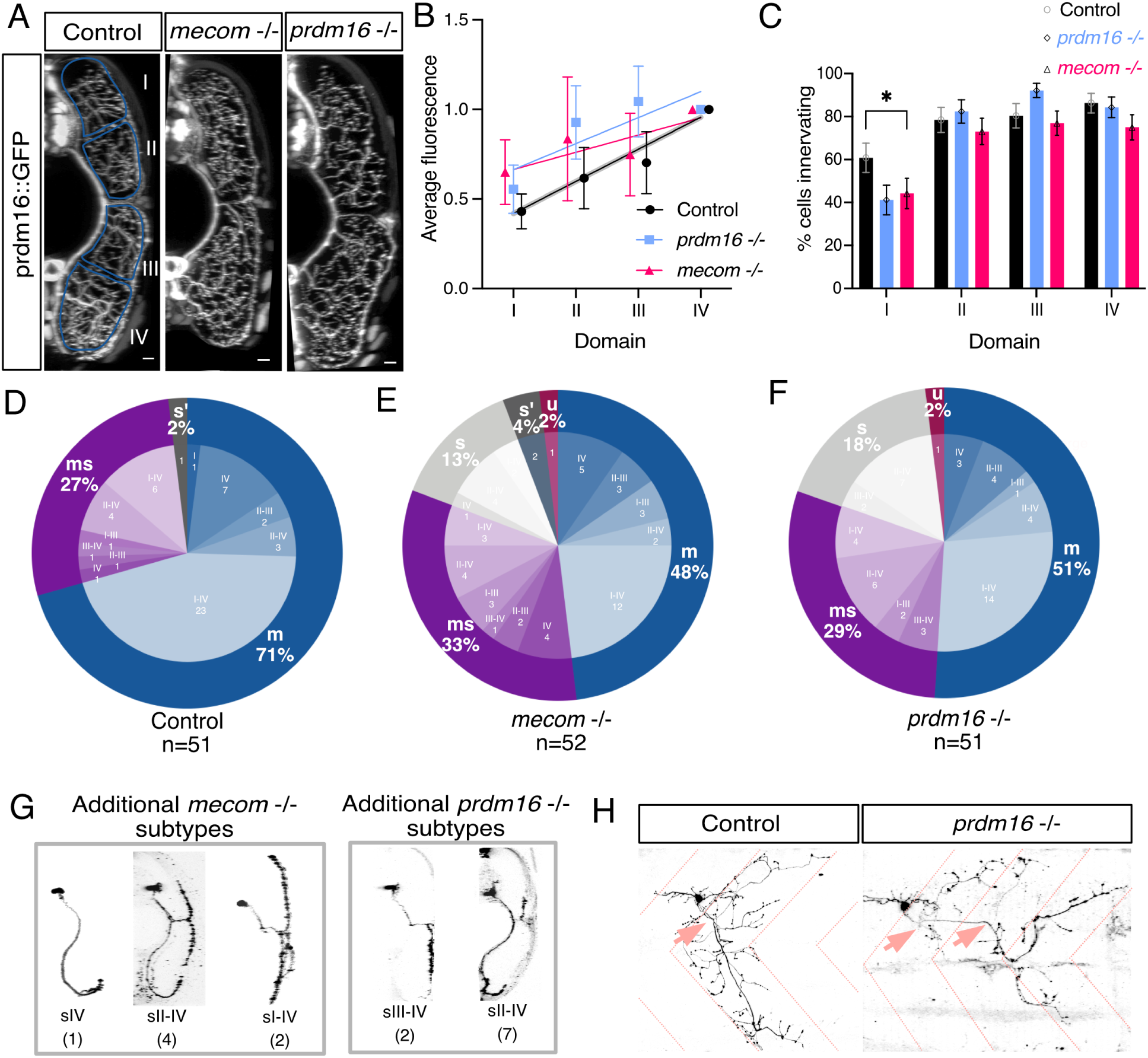
*prdm16* and *mecom* are required for normal muscle innervation. (A) Transverse view of the zebrafish tail showing Prdm16^+^ MN muscle innervation in *Tg(prdm16::GFP)* control, *mecom* mutant, and *prdm16* mutant larvae at 5 dpf. Scale bars, 10 µm. (B) Average GFP fluorescence of each muscle quadrant normalized to the fourth quadrant for control *Tg(prdm16::GFP)* fish (n = 6), *mecom* mutant (n = 9), and *prdm16* mutant (n = 10) larvae at 5 dpf. Error bars indicate SEM. Lines illustrate linear regression for each condition and grey area surrounding control indicates standard deviation of the slope, ±0.00555. (C) Percentage of cells from control, *mecom* mutant, and *prdm16* mutant *Tg(prdm16:GAL4)* sparse label experiments in Figures D-F that innervated each dorsoventral muscle quadrant. Asterisk indicates significant difference (*prdm16 -/-* domain I: p=0.000891, *mecom -/-* domain I: p=0.004121), all other domains were not significantly different (p > 0.00625). (D) Pie chart showing subtype proportions labeled in *Tg(prdm16:GAL4)* sparse label for control larvae. Replicated from Figure 5E. (E) Pie chart showing subtype proportions labeled in *Tg(prdm16:GAL4)* sparse label for *mecom* mutant larvae. (F) Pie chart showing subtype proportions labeled in *Tg(prdm16:GAL4)* sparse label for *prdm16* mutant larvae. (G) Example images of all sparsely labeled subtypes unique to mutants (E & F). Number in parentheses indicates number of times subtype was labeled. (H) Example of an unclassifiable cell found in mutant sparse label experiment from the lateral view. Dashed coral lines indicate myotome boundaries. Coral arrows indicate motor exit points. Controls can indicate either WT or *prdm16* +/- or *mecom* +/- animals which are phenotypically indistinguishable.

To further explore a potential change in muscle innervation, we performed sparse labeling by injecting a *pUAS:DsRed* plasmid in a *Tg(prdm16:GAL4)* mutant background. Consistent with the overall fluorescence quantification, we saw a significant decrease in increase in the proportion of cells in mutants that innervate quadrant I (Figure 6C, *prdm16-/-* p=0.000891, *mecom -/-* domain I: p=0.004121), all other domains p > 0.00625).

More strikingly, in contrast to control larvae, we observed that 13% (7/52) of labeled cells in *mecom* mutants and 18% (9/51) of labeled cells in *prdm16* mutants were s-type with innervation of only slow muscle (Figures 6E, 6G, S9A and S9B, Table 1). Hypothesis testing by resampling (Methods) allowed us to reject the null that the cell types we observed in mutants reflect the wild-type Prdm16^+^ distribution of cells: there were fewer m types (p = 0.0034, *prdm16* mutants, p = 0.0011 *mecom* mutants); more s types (p = 0, *prdm16* mutants, p = 0.0001, *mecom* mutants); and comparable numbers of ms types (p = 0.392, *prdm16* mutants, p = 0.1786, *mecom* mutants). In addition, in both *prdm16* and *mecom* mutants, we observed MNs with axons exiting from multiple motor exit points and extending beyond the normal myotome boundaries. We considered these “unclassifiable” or “u” types (Figures 6E, 6F and 6H). While the high mortality rate of double mutants prohibited sparse labeling, we were able to assess *prdm16*-/-;*mecom*+/- larvae. Compared to single mutants, we observed nearly twice as many s-type MNs at 31% (10/32) and a higher proportion of unclassifiable MNs at 9% (3/32) (Figure S8K, Table 1). Compared to the wild-type Prdm-16^+^ distribution, there were fewer m types (p = 0), more s types (p = 0), and comparable numbers of ms types (p = 0.5274).

To determine if *prdm16* or *mecom* mutants had swim speed deficits, we utilized a previously established free-swimming analysis pipeline to compare mutant larvae and swim bladder-less controls ^51^. *mecom* mutants were more likely to execute slower and shorter bouts and have an exaggerated nose-up posture compared to swim bladder-less controls (Movie 1, Figure S8L). *prdm16* mutants had more drastic posture and coordination defects compared to mecom mutants, prohibiting quantitative analyses of swimming behaviors (Movie 2). These speed changes and instability could be from a miscoordination of motor programs or abnormal fast MN development and activity, but the global nature of the mutations makes it impossible to attribute these effects specifically to fast MN changes.

### Molecular changes in fast motor neurons with the loss of Prdm16 or Mecom

To investigate molecular changes in fast MNs after loss of *prdm16* or *mecom*, we compared the transcriptional profiles of control and single mutant larvae MNs. We first sorted *Tg(Prdm16::GFP)*;*Tg(olig2:DsRed2)* mutant embryos at 5 dpf by the absence of a swim bladder (Figure S10A). We isolated MNs by FACS for both GFP and DsRed to allow the enrichment and identification of putative fast MN clusters (Figure S10E). The dim green fluorescence at 5 dpf and bleedthrough from the red channel limited our ability to separate GFP^+^ and dsRed^+^ MNs. Combined, the two samples for each condition were 14,882 cells for WT, 9,005 cells for *mecom* mutants, and 12,445 cells for *prdm16* mutants. The overall quality of the samples was comparable (Figure S10C). Cells from all samples were pooled for downstream analysis. MN clusters were identified using *chata* expression (18,848 MNs total; Figures S10B and S10D) and reclustered for further analysis (Figure 7A). Since *prdm16* and *mecom* are no longer expressed in fast MNs at 5 dpf, we identified the fast MN cluster (14) in the WT sample using other markers including *calca*, *calb2b*, and *pcp4a*. The limited representation of fast MNs at 5 dpf is likely due to a decay of fluorescence in older MNs.

**Figure 7:**
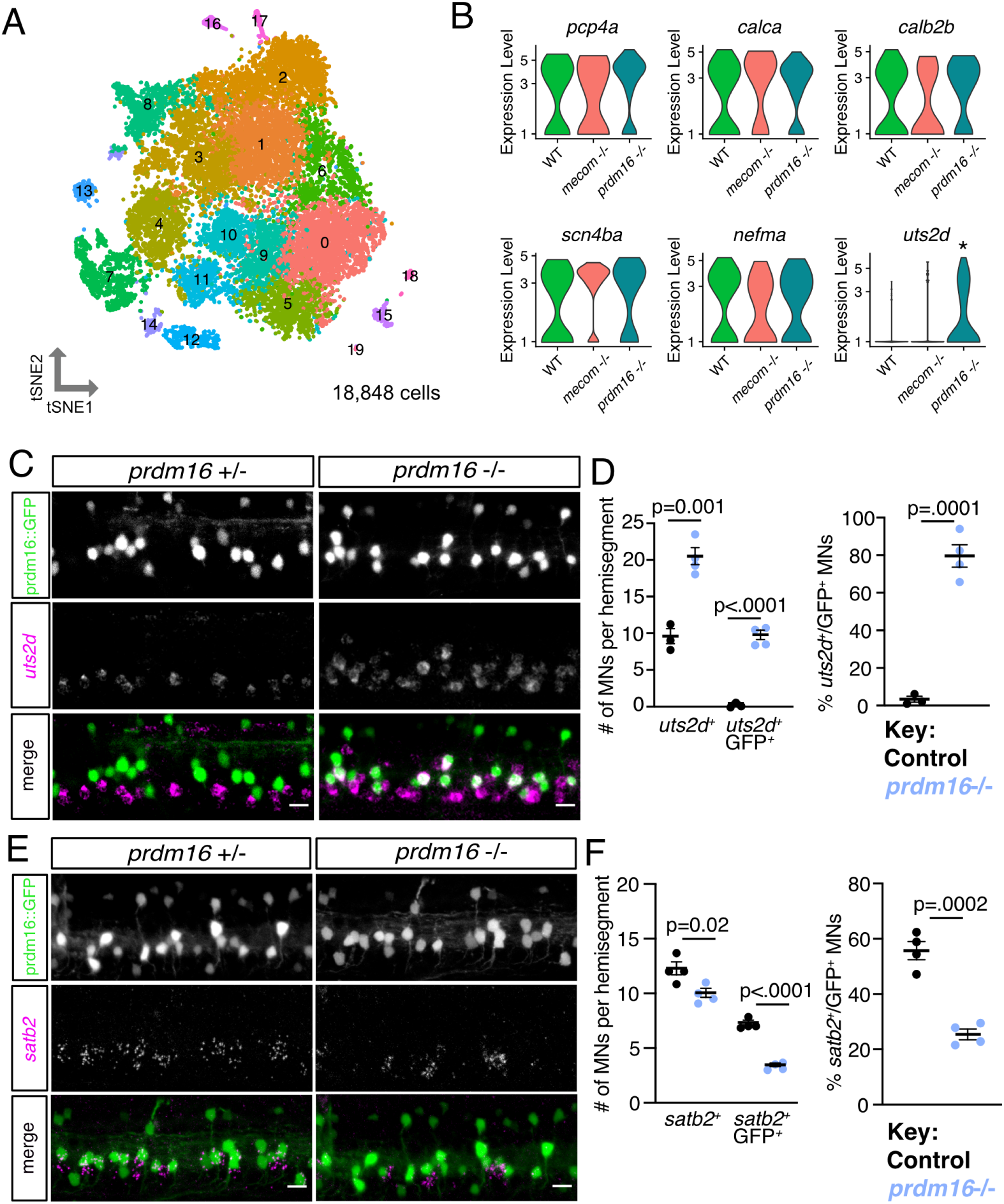
Erosion of fast motor neuron transcriptional profiles in *prdm16* mutants. (A) tSNE representing all MNs from WT (8145), *mecom* mutant (4210), and *prdm16* mutant (6493) larvae at 5 dpf. Cluster 14 was determined to be fast MNs (WT = 88, *mecom* mutant = 40, and *prdm16* mutant = 65). (B) Violin plots representing log fold expression of fast MN markers (*pcp4a*, *calca*, *calb2b*, *scn4ba*, *nefma*) and slow MN marker (*uts2d*) in cluster 14 of (A) identified as normally Prdm16^+^ fast MNs. *uts2d* showed a three log fold change increase in *prdm16* mutants (p adj = 0.0194). (C) FISH of *uts2d* in control and *prdm16* mutant larvae at 5 dpf. Scale bars, 10 µm. (D) Left: Quantification of *uts2d*^+^ cells and GFP^+^ cells in *Tg(prdm16::GFP)*; *uts2d*^+^ cells at 5 dpf as in C. Right: Percentage of total GFP^+^ cells that express *uts2d*. Line indicates average with SEM. Unpaired t-tests. (E) FISH of *satb2* in control and *prdm16* mutant larvae at 5 dpf. Scale bars, 10 µm. (F) Left: Quantification of *satb2*^+^ cells and GFP^+^; *satb2*+ cells at 5 dpf as in E. Right: Percentage of total GFP^+^ cells that express *satb2*. Line indicates average with SEM. Unpaired t-tests.

We then performed differential gene expression analysis between each mutant and the WT sample within fast MN cluster 14. The gene with the largest fold change between WT and mutant samples was *uts2d* which had a three log-fold increase in *prdm16* mutants (p adj = 0.0194; Figure 7B). Analyses by FISH in control embryos at 5 dpf showed that *uts2d* is normally expressed within slow MNs ventral to Prdm16^+^ MNs (Figures 7C and 7D). In *prdm16* mutants, the majority of Prdm16^+^ MNs (80%) ectopically expressed *uts2d* (Figure 7D). To determine if fast MN molecular signatures were disrupted within the mutant samples, we also screened additional genes normally expressed within Prdm16+ fast MNs. There was no significant change in expression of *pcp4a*, *calca*, *calb2b*, *scn4ba*, or *nefma* within *prdm16* or *mecom* mutant cluster 14 (Figure 7B). While we confirmed no loss in *pcp4a* or *calb2b* expression by FISH, we did find a partial loss of *satb2* expression in *prdm16* mutant fast MNs (Figures 7E and 7F). The limited detection of changed genes in mutants may be due to the small number of fast MNs in the 5 dpf dataset.The gain of expression of *uts2d* and partial loss of *satb2* expression suggests the loss of *prdm16* alters the transcriptional profiles of fast MNs to be more slow-like.

## DISCUSSION

In this study, we show that axial MN subtypes are remarkably diverse and identify transcriptional determinants of MN functional subtypes. Leveraging the larval zebrafish spinal cord, we defined both conserved and novel markers that define functional subtypes both during early development and at functional maturity. We found that two conserved transcription factors – Prdm16 and Mecom – are expressed in fast secondary MNs. Loss of these genes leads fast secondary MNs to acquire features that normally define slow MNs. These data provide novel insights into molecular diversity of vertebrate spinal MNs, and establish that MN functional subtypes are determined by intrinsic genetic programs acting in early development.

### Molecular diversity of axial motor neurons

Our findings reveal unappreciated transcriptional diversity in axial MNs that offers insight into the molecular underpinnings of MN specification. We observed that zebrafish fin and tetrapod limb MNs (1) have conserved expression of *foxp1* and *lhx1a* and (2) both lack expression of *lhx3*/*lhx4* ^44, 45^. Interestingly, while *lhx3/4* were excluded from fin MN clusters, they were widely expressed within zebrafish axial MNs. In mammals, the specification of ventrally-projecting MN subtypes requires downregulation of *lhx3/4*, as sustained expression of *lhx3* in all MNs blocks the specification of limb, preganglionic and hypaxial subtypes ^52, 53^. One interpretation of these findings could be that Lhx3^+^ MNs represent an ancient and more molecularly simple MN subtype fate ^54^.

We show that axial MNs are highly diverse, and this molecular heterogeneity exists across vertebrate species. Two new markers, *prdm16* and *mecom*, as well as two recently identified ^48^ MMC subtype markers, *satb2* and *nr2f2*, mark subpopulations of zebrafish axial MNs. *satb2* and *nr2f2* appear to mark two populations of secondary MNs with *satb2* expressed in a population overlapping with *prdm16*+^+^ fast MNs and *nr2f2* labeling putative slow MNs and progenitors. We conclude that MN subtype markers, including those marking axial motor columns and their diverse subpopulations of pools, are conserved among vertebrates.

Our finding that molecular signature of MN subtypes persist from early development to functional maturity extends previous work towards a more complete description of zebrafish axial MNs. Previously, relating the well-established anatomical and functional diversity of zebrafish axial MNs to molecular heterogeneity was an open challenge. For example, the developmental timing of expression of Islet, Mnx, and Lhx family members are known to be important for specifying features of primary MNs, but these factors are also dynamically expressed within secondary MN subtypes ^16–19, 21, 22^. Our findings indicate that both slow and fast secondary MNs express selective markers of subtype identity. Importantly, while we classified dorsally positioned markers as “fast” and ventrally positioned ones as “slow,” prior work further refines these categories, including primary/secondary and intermediate MN distinctions. The absence of intermediate muscle or defined intermediate subtypes in the larval zebrafish underscores the importance of performing similar functional classification with accompanying molecular profiling in juvenile / adult zebrafish ^28, 30^. We hypothesize that MNs located near the boundary of the dorsal/ventral (fast/slow) distinction that project along both the medial and septal nerves, and/or those that express genes that mark a portion of the fast or slow groups are plausible candidates.

### Prdm16 and Mecom specify fast motor neuron subtypes

Our loss-of-function experiments establish two transcription factors, Prdm16 and Mecom, as indispensable in the development of a subset of fast MNs. Prdm16 and Mecom exclusively mark secondary MNs that innervate fast muscle. Loss of either *prdm16* or *mecom* leads to fast MN developmental deficits including the acquisition of anatomical and molecular features of slow subtypes. In both *prdm16* and *mecom* mutants, fast MNs retain many of their molecularly defined features, but now express a marker normally restricted to slow MNs. The absence of a complete fast-to-slow fate transformation could be due to functional compensation between these factors, but the high lethality rate of double mutants precludes a detailed molecular characterization. Alternatively, these factors may play more specific roles, by acting in fast MN to repress subsets of genes that define slow MN fate.

Both Prdm16 and Mecom have been shown to function as repressors in certain contexts ^55^. Our findings argue against a role for *prdm16* and *mecom* in neurogenesis as both genes are expressed post-mitotically and presumptive fast MNs are still generated in mutant lines, although they acquire molecular features of slow MNs. We propose that they work to suppress gene programs that define alternate MN subtype fates. Such suppression is consistent with our observation that *prdm16* mutants derepress *uts2d*, a gene of unknown function ^47^ whose homolog *uts2b* is expressed in mammalian axial MNs ^56^.

Prior to our discovery of Prdm16 and Mecom as a determinant of fast secondary MNs, the mechanisms for functional subtype specification remained unresolved. One proposal was that MN functional subtype identity is plastic and sensitive to retrograde influences from muscle fibers once the targets are reached ^57–60^. Alternatively, differences between MN functional subtypes might reflect early-defined intrinsic programs ^42, 61–63^. Our results provide strong evidence that an intrinsic transcription factor code determines functional subtype and is supported by our finding that expression of Prdm16 and Mecom in MNs precedes secondary MN muscle innervation (data not shown) ^64^. Broadly, functional subtype specification of MNs therefore resembles the transcriptional codes that specify classes of functionally distinct subtypes elsewhere in the nervous system ^65–69^.

### Implications for locomotor speed circuit organization and assembly

Our findings establish that functional subtype-restricted molecular codes exist in conjunction with, and likely control, established temporal aspects of locomotor speed circuit maturation. To regulate speed, zebrafish rely on circuits comprised of descending neurons, spinal interneurons, and MNs ^27, 28, 70, 71^. Shared functional properties (fast/slow) among cell types within speed control circuits reflect birthdate ^27, 37, 72, 73^, similar to other sensorimotor circuits ^74–78^. Spatial and temporal morphogen gradients, as well as sequential expression of transcription factors, have been implicated in generating spinal neuron subtype diversity across species ^79, 80^. Birthdate-dependent subtype differences have been linked to temporally regulated post-mitotic transcription factor programs in vertebrates ^81, 82^ and invertebrates ^83, 84^. We propose that Prdm16 and Mecom operate as intrinsic determinants acting within a temporal program that defines MN functional subtype fates.

Broadly, our work defines novel molecular signatures for diverse spinal MN subtypes and reveal genetic mechanisms of MN functional subtype specification in the vertebrate nervous system. Every vertebrate muscle is innervated by both slow and fast MN subtypes; the discovery that a transcription factor code establishes functional subtype identity thus speaks broadly to key mechanisms of motor control. Further, these data establish key regulatory nodes in fast and slow MN development, potentially opening new avenues to understand the differential susceptibility of MNs in diseases such as ALS. Finally, identifying the hallmarks of functional subtype determination permit comparative studies of molecular underpinnings of locomotor control and its evolutionary origins.

## MATERIALS AND METHODS

### Fish Care

All procedures involving zebrafish larvae (*Danio rerio*) were approved by the Institutional Animal Care and Use Committee of the New York University Grossman School of Medicine. Fertilized eggs were collected and maintained at 28.5°C on a standard 14/10 hour light/dark cycle. Before 5 dpf, larvae were maintained at densities of 20-50 larvae per petri dish of 10 cm diameter, filled with 25-40 mL E3 with 0.5 ppm methylene blue.

### Mouse/chick care

All procedures involving mice or chicks were performed in accordance with the US National Institutes of Health Animal Protection Guidelines and approved by the Institutional Animal Care and Use Committee of the New York University Grossman School of Medicine. Animal ages are indicated in the main text and figure legends. No sex-specific differences in reported phenotypes are expected, but were not formally tested.

### Skate care

*Leucoraja erinacea* embryos of the desired stages were obtained from the Marine Resources Department of Woods Hole Marine Biology Laboratory (MBL). Animals at MBL are maintained in accordance with procedures approved by the MBL Institutional Animal Care and Use Committee following standards established by the National Institutes of Health. Prior to use, egg cases were maintained in reconstituted Instant Ocean (Aquarium Systems) at 16°C with a standard aquarium filtration system. Before manipulation, embryos were anesthetized with MS-222 (0.17 g/L; Sigma-Aldrich) at room temperature for 10 min ^85^.

### Fish lines

Experiments were done on the wildtype or the *mitfa*-/- background to remove pigment. We used *Tg(olig2:DsRed2)^vu19 38^*. *Tg(olig2::GFP)* line is *Et(-0.6hsp70l:GAL4-VP16)^s1020t^* ^86^ crossed with *Tg(5xUAS:EGFP)* ^87^. *Tg(prdm16:GAL4)* is gSAIzGFFM1116A [Kawakami lab, insertion site CACACTTCAGT-GCTTTTGCAATTTTCTAGTCTGCATCAACTGCTCTGAGCT located upstream of the prdm16 locus on Chr 8 in GRCz10].

*Tg(prdm16::GFP)* is *Tg(prdm16:GAL4)* crossed with *Tg(5xUAS:EGFP)*.

### Generation of mutants

*prdm16* and *mecom* mutant lines were generated via CRISPR-based mutagenesis. Guide RNAs (gRNAs) were designed using the CHOPCHOP online tool ^88^. Location of guides was chosen to be in the largest exon based on its presence in multiple described isoforms of each gene. gRNAs were incubated with Cas9 protein before co-injection into embryos at the single cell stage. Injected embryos were raised and their progeny were screened by genotyping to identify germline mutations. Multiple founders were identified and their progeny with variable mutations consistently lacked swim bladders. We focused on one of these alleles for the following experiments. Each mutant had a sizable deletion (*mecom^nyc41^* mutant = 25 bp, *prdm16^nyc142^* mutant = 17 bp, *Tg(prdm16:GAL4);prdm^nyc587^* mutant = 22 bp) detectable by genotyping that created a nonsense mutation and a premature stop codon. Loss of protein in mutants were validated using antibodies.

### Dissection, cell dissociation, and FACS

DsRed positive cells were isolated from 2 dpf or 5 dpf control, *mecom* mutant, or *prdm16* mutant zebrafish embryos on an *Tg(olig2:DsRed2)* or *Tg(prdm16::GFP)*;*Tg(olig2:DsRed2)* background. Trunk and tail tissue were separated from cranial tissue. Tissue was then finely chopped using a razor blade, dissociated using papain, filtered, and resuspended for sorting. Cells were sorted using the Sony SH800 FACS Cell Sorter. GFP^-^, DsRed2^-^, and single fluorophore control embryos were also included as controls for the FACS setup. DAPI was used for a live/dead marker. Sorted cells were then counted using a hemocytometer and spun down to resuspend at a higher concentration. Cells were then processed using the standard 10x Genomics and CellRanger(v5.0.1) pipeline ^89^. Raw sequencing reads were mapped to the zebrafish reference genome (build GRCz11).

### Single cell analysis

To evaluate the 2 dpf scRNAseq data, we used Seurat v4^90^. To ensure only high-quality cells were retained for downstream analysis, cells with fewer than 200 genes, more than 5,000 genes, over 6% mitochondrial counts, and total RNA counts over 30,000 were removed. Data was normalized using the NormalizeData Seurat function and variable features were identified using FindVariableFeatures. The top 2000 variable genes were used in the PCA. Based on Jack Straw and an elbow plot of 30 principal components ^91^, 21 principal components were used in the downstream analysis. For the 5 dpf data, we used a *Tg(prdm16::GFP)*;*Tg(olig2:DsRed2)* transgenic line. To ensure only high-quality cells were retained for downstream analysis, cells with fewer than 200 genes, more than 3,500 genes, over 10% mitochondrial counts, and total RNA counts over 30,000 were removed. Since samples were sorted for both DsRed and GFP, we used the top 2000 variable genes to find anchors to integrate the GFP^+^ and GFP^-^ negative populations using IntegrateData. This was also done to combine the WT, *mecom* mutant, and *prdm16* mutant datasets. There, the top 2000 variable genes were used in the PCA, and 21 principal components were used in the downstream analysis. To isolate MN clusters, we reran the above analyses on data from clusters that expressed *chata*. After MNs were reclustered, we re-evaluated mitochondrial and gene counts within each cluster. In the 2 dpf data, three clusters were identified as having high mitochondrial counts and low gene counts. These were determined as outliers within the data. After their removal, the remaining cells were reclustered. For 2 dpf data, 20 principal components were used for MN clustering. For 5 dpf data, 30 principal components were used for MN clustering. For the 2 dpf v 5 dpf analysis, R package CIDER (v0.99.0) ^92^ was used to assess inter-group similarity between 2 dpf and 5 dpf clusters . Gene Ontology analysis was done using ShinyGO (v0.76.3) ^93^.

### RNA Velocity

The .loom file from velocyto (v0.17.17) ^40^ was used to calculate splicing information for each gene, which was then used to compute RNA velocities for each cell according to standard parameters in scVelo (v0.2.4) ^41^. UMAP coordinates calculated with Seurat were exported and used at the basis for the velocity streamline plot in Figure 1E. Pseudotime was then computed based on RNA velocity information according to suggested parameters in the scVelo package.

### Integration with Scott et. al. 2021

Raw .fastq files were downloaded from 1 day and 1.5day timepoints published in ^39^, 2021 NCBI GEO #GSE173350. For this analysis, the .fastq files for the raw data for both Scott et al. and our datasets were analyzed via CellRanger v3.0.2 as done in their study. We used the .fastq files to produce .loom files (with splicing data for each gene) via velocyto splice calling. We then processed these .loom files for RNA velocity as above, and then we integrated both published datasets with our later timepoints using HarmonyPy (v0.0.5) ^94^.

### Mouse *in situ* hybridization

Embryos were fixed in 4% PFA for 1.5-2 hr at 4°C. Embryos were washed 5-6 times in cold PBS, 15-30 min for each wash, and incubated overnight in 30% sucrose. Antisense riboprobes were generated using the Digoxigenin-dUTP (SP6/T7) labeling kit (Sigma-Aldrich). RNA was extracted from E12.5 embryos using TRIzol (Invitrogen) and cDNA was generated using SuperScript II Reverse Transcriptase (Invitrogen). Riboprobes were prepared by PCR using cDNA templates, incorporating a T7 polymerase promoter sequence in the antisense oligo. Sections were dried for 10-15 min at room temperature, placed in 4%PFA, and fixed for 10 min at room temperature. Slides were then washed three times for 3 min each in PBS, and then placed in Proteinase K solution (1 mg/ml) for 5 min at room temperature. After an additional PFA fixation and washing step, slides were treated in triethanolamine for 10 min, to block positive charges in tissue. Slides were then washed three times in PBS and blocked for 2-3 hr in hybridization solution (50% formamide, 5X SSC, 5X Denhardt’s solution, 0.2 mg/ml yeast RNA, 0.1 mg/ml salmon sperm DNA). Prehybridization solution was removed, and replaced with 100 mL of hybridization solution containing 100 ng of DIG-labeled antisense probe. Slides were then incubated overnight (12-16 hr) at 72°C. After hybridization, slides were transferred to a container with 400 mL of 5X SSC and incubated at 72°C for 20 min. During this step, coverslips were removed using forceps. Slides were then washed in 400 mL of 0.2X SSC for 1 hr at 72°C. Slides were transferred to buffer B1 (0.1M Tris pH 7.5, 150mM NaCl) and incubated for 5 min at room temperature. Slides were then transferred to staining trays and blocked in 0.75 ml/slide of B1 containing 10% heat inactivated goat serum. The blocking solution was removed and replaced with antibody solution containing 1% heat inactivated goat serum and a 1:5000 dilution of anti-DIG-AP antibody (Roche). Slides were then incubated overnight at 4°C in a humidified chamber. The following day, slides were washed 3 times, 5 min each, with 0.75 ml/slide of buffer B1. Slides were then transferred to buffer B3 (0.1 M Tris pH 9.5, 100 mM NaCl, 50 mM MgCl2) and incubated for 5 min. Slides were then developed in 0.75 ml/slide of B3 solution containing 3.5 ml/ml BCIP and 3.5 ml/ml NBT for 12-48 hr. After color development, slides were washed in ddH20 and coverslipped in Glycergel (Agilent). A more detailed *in situ* hybridization protocol is available on the Dasen lab website: https://www.dasenlab.com/lab-protocols.

### Zebrafish *in situ* hybridization (ISH)

Preparation of RNA probes was done as described above. DNA templates were made using a zebrafish 30 hpf cDNA library generated from polyA RNA extracted with Trizol. Whole mount *in situ* hybridization was performed as previously described ^95^. Embryos were fixed in 4% PFA overnight at room temperature. Following fixation, they were permeabilized in methanol at 20°C, rehydrated, permeabilized with 10mg/ml proteinase K in PBST for 8min at room temperature, and post-fixed in 4% PFA for 20min at room temperature. Embryos were bleached in a 10% H_2_O_2_, 10% KOH solution prepared in ddH_2_O and washed twice for five minutes in PBS with 0.1% Tween (PBST). Blocking and probe hybridization were performed at 68°C overnight. Following probe hybridization and washes, embryos were blocked in 2%BSA in PBST and incubated with anti-DIG-AP antibody overnight at 4°C. Embryos were washed and developed in NBT/BCIP solution overnight at room temperature.

### Zebrafish fluorescent *in situ* hybridization (FISH)

Protocol adapted from ^96^: Hybridization chain reaction (HCR) probes were generated by the HCR 3.0 probe maker ^97^ based on methodology generated by ^98^. Probe pairs were created based off the sense sequence of the gene cDNA from NCBI. Embryos were previously fixed at the desired age with 4% PFA in PBS overnight at 4°C. Following fixation, the embryos were dehydrated in 100% methanol at -20 °C until used. When needed, the embryos were rehydrated through a titer of methanol and PBST, washed two times with PBST for 5 minutes, and permeabilized in pre-chilled acetone at -20°C for 12 minutes. After acetylation, the samples underwent three PBST washes for five minutes. Further permeabilization was performed by placing the embryos in 10 mg/ml proteinase K in PBST at room temperature. Optimal proteinase K incubation time was determined by embryo age. Two day post fertilization embryos were digested for 20 minutes and five day post fertilization embryos for 50 minutes. The samples were then rinsed with PBST three times for five minutes and post-fixed in 4% PFA in PBST for 20 minutes at room temperature. Embryos were washed 5 times with PBST for 5 minutes each and incubated in pre-warmed whole mount probe hybridization buffer (from Molecular Instruments) at 37°C for 30 minutes. Incubation with HCR probe mixture solution was performed overnight at 37°C. Four 15 minute washes of preheated probe wash buffer (from Molecular Instruments) were performed at 37°C. The embryos were then moved to room temperature, washed two times with 5x SSCT for five minutes each, and placed in the probe amplification buffer for 30 minutes. As the larvae underwent preamplification, aliquots of amplifier hairpins were snap cooled by heating to 95°C for 90 seconds and cooling at room temperature in the dark for 30 minutes. The embryos were incubated in the dark with the amplification buffer and snap cooled hairpins overnight at room temperature. Excess amplifier hairpins were removed by washing with two five minutes washes, two 30 minute washes, and one last five minute wash of 5x SSCT. Finally, they were washed three times with PBST for five minutes each.

### Chick/skate immunohistochemistry

For antibody staining of sections, slides were first placed in PBS for 5 min to remove OCT. Sections were then transferred to humidified trays and blocked for 20-30 min in 0.75 ml/slide of PBT (PBS with 0.1% Triton) containing 1% Bovine serum albumin (BSA). The blocking solution was replaced with primary staining solution containing antibodies diluted in PBT with 0.1% BSA. Primary antibody staining was performed overnight at 4°C. Slides were then washed three times for 5 min each in PBT. Fluorophore-conjugated secondary antibodies were diluted 1:500-1:1000 in PBT and filtered through a 0.2 mm syringe filter. Secondary antibody solution was added to slides (0.75 ml/slide) and incubated at room temperature for 1 hour. Slides were washed three times in PBT, followed by a final wash in PBS. Coverslips were placed on slides using 110 mL of Vectashield (Vector Laboratories).

### Zebrafish immunohistochemistry

All steps were done in a 1.5 mL Eppendorf tube. Dechorionated embryos were fixed in 8% PFA in 2X Fix Buffer (8.0 g sucrose + 30 µl 1M CaCl_2_ + 10 ml 10X PBS in 80 µl ddH_2_O) at 4°C for two hours. They were then rinsed once and washed three times for 20 minutes each in PBS with 0.1% Tween (PBST). Embryos were put in 150 mM Tris for 5 min at RT, then put in a 70° water bath for 15 min. Embryos were then washed twice for 5 minutes in PBST. Embryos were then put in 0.5% Triton in PBS for 15 minutes. Embryos were then bleached only if pigmented in a 10% H_2_O_2_, 10% KOH solution prepared in ddH_2_O and washed twice for five minutes in PBS with 0.1% Tween. Embryos were blocked for one hour in PBT, 2.5% DMSO, and 5% Heat Inactivated Goat Serum. Primary antibodies were diluted in PBT with 1% BSA, and embryos were incubated at 4°C for 3 or 4 nights. Embryos were then rinsed once and washed four times for 30 minutes each in PBS with 0.1% Tween. Secondary antibodies were diluted in PBT with 1% BSA, and embryos were incubated at 4°C overnight. Embryos were then rinsed once and washed four times for 30 minutes each in PBST. Embryos were mounted in agarose or on a slide in Vectashield (Vector Laboratories) to be imaged. Protocol adapted from ^16^.

### Sparse labeling

For sparse labeling, the GAL4-UAS system was used to drive mosaic expression of a reporter construct with a membrane-targeted fluorophore selectively in Prdm16-expressing neurons. Mosaic expression was obtained by injecting 40 ng/µl of a *pUAS:DsRed* plasmid with or without *Tol2* messenger RNA into oneto two-cell *Tg(prdm16:GAL4)* embryos using a microinjector. *pUAS:DsRed* is 5UAS zfSynaptophysin:GFP-5UAS DsRedExpress [RRID:Addgene_74317] originally from ^50^.

### Imaging

Live zebrafish were anesthetized in 0.02% MS-222 and mounted on their side in 2% low-melt agarose for image collection. Larvae were imaged between 2-5 dpf on a Zeiss LSM800 confocal microscope with a 20x water-immersion objective (Zeiss W Plan-Apochromat 20x/1.0 DIC CG=0.17 M27 75mm). For single labeled neurons, a z-stack was acquired to capture the entire morphology of the cell using the 20x objective with an optical section of 1.5 µm. All images were taken within the mid-trunk of the embryos or larvae (between segments 6 and 20). All lateral images are maximum intensity projections through half of the spinal cord and cropped to two hemisegments. All transverse images were created by rotating and interpolating the z-stack 90° in Fiji ^99^. The spinal cord transverse images are maximum projections of half of a hemisegment. The muscle transverse images are cropped according to the myotome boundary prior to rotation, and a maximum projection created from the entire myotome. Due to the dim fluorophore expression within older MN populations (because of the progenitor promotor), gamma was adjusted to visualize the entire population evenly (≥0.45).

### Sparse label MN subtype classification

Only hemisegments containing one labelled MN were used for subtype classification based on axon morphology. Three-dimensional images were reconstructed and analyzed using Fiji as described above ^99^.

### Antibodies

Antibodies against LIM HD proteins and other proteins were used as described ^20, 21, 100^. Additional antibodies were obtained as follows: rabbit anti- Evi1/Mecom (Cell Signaling, RRID: AB_2184098); chicken and rabbit anti-GFP (Invitrogen, RRID: AB_2534023). Guinea pig anti-zebrafish Prdm16/Mecom antibody and Rabbit anti-zebrafish Mecom antibody were made by Covance (Denver, PA) with custom chosen peptides. Serial dilutions were done to test optimal concentrations to use for zebrafish whole mount staining. Prdm16/Mecom antibody was found to also work in chick and mouse spinal cord.

### Mapping soma positions

Threedimensional images were reconstructed and analyzed using Imaris (Bitplane). Neural soma positions were first auto-determined using the Spots function, then verified by eye to exclude any non-soma structures that were detected. Coordinates were then exported from Imaris and plotted using custom MATLAB scripts based on previously published scripts ^73, 101^. To account for variations in spinal cord height and width across different fish, coordinates were normalized relative to the dorsal-most and ventral-most aspects of the spinal cord as well as along mediolateral landmarks marked medially by center of the spinal cord, and laterally by the lateral-most boundary between the spinal cord and the axial musculature. The dorsal and ventral aspects of the spinal cord were determined from differential interference contrast (DIC) images captured during image acquisition on the confocal microscope. Variations in myotome width were also accounted for by normalizing coordinates relative to the most rostral and caudal planes of the segment marked by the ventral root exit point from the hemisegment before as the most rostral and the ventral root of the neuron’s segment as the most caudal. Normalized coordinates from all MNs or interneurons were then either plotted as a scatter plot or pooled to generate separate contour plots that represent the variation in densities. Histograms were also generated to summaries the positional variability on the x and y axes of the scatterplots. Distribution contours were made in MATLAB using the kde2d function (MATLAB File Exchange), which estimates a bivariate kernel density over the set of normalized coordinates using a 32x32 grid ^102, 103^. Contour plots were then generated from the calculated densities using the contour function in MATLAB. Contour lines are displayed in increasing increments of one-sixth of the peak density where the innermost contour represents the highest density of coordinates.

### Behavioral measurement

All behavioral experiments were performed at 5 dpf. Control or mutant larvae were put into an air-tight glass cuvettes (93/G/10 55x55x10 mm, Starna Cells, Inc., Atascadero, CA, USA) at 5 dpf sealed with parafilm and a custom fit SYLGARD cap. All groups of fish were checked for swim bladders following the experiment and validated by genotyping. If any fish inflated their swim bladder or was of the wrong genotype, the entire group’s data was thrown out. Larvae were filmed in groups of 8 siblings in a glass cuvette filled with 24-26 mL E3 and recorded for 24 hours as described previously ^51, 104^. The larvae’s normal 14/10 light/dark cycle was maintained within the light-tight box using LED lights on a timer. Video was captured using digital CMOS cameras (Blackfly PGE-23S6M, FLIR Systems, Goleta CA) equipped with close-focusing, manual zoom lenses (18-108 mm Macro Zoom 7000 Lens, Navitar, Inc., Rochester, NY, USA) with f-stop set to 16 to maximize depth of focus. The field-of-view, approximately 2x2 cm^2^, was aligned concentrically with the tank face. A 5W 940nm infrared LED back-light (eBay) was transmitted through an aspheric condenser lens with a diffuser (ACL5040-DG15-B, ThorLabs, NJ). An infrared filter (43-953, Edmund Optics, NJ) was placed in the light path before the imaging lens. Digital video was recorded at 40 Hz with an exposure time of 1 ms. Kinematic data was extracted in real time using the NI-IMAQ vision acquisition environment of LabVIEW (National Instruments Corporation, Austin, TX, USA). Background images were subtracted from live video, intensity thresholding and particle detection were applied, and age-specific exclusion criteria for particle maximum. Feret diameter (the greatest distance between two parallel planes restricting the particle) was used to identify larvae in each image. In each frame, the position of the visual center of mass and posture (body orientation in the pitch, or nose-up/down, axis) was collected. Posture was defined as the orientation, relative to horizontal, of the line passing through the visual centroid that minimizes the visual moment of inertia. A larva with posture zero at any given time has its longitudinal axis horizontal, while +90° is nose-up vertical, and -90° is nose-down vertical.

### Behavioral analysis

Data analysis and modeling were performed using Python 3. As previously described ^51, 104^ epochs of consecutively saved frames lasting at least 2.5 sec were incorporated in subsequent analyses if they contained only one larva. Instantaneous differences of body particle centroid position were used to calculate speed. Inter-bout intervals (IBIs) were calculated from bout onset times (speed positively crossing 4 mm/sec) in epochs containing multiple bouts, and consecutively detected bouts faster than 10 Hz were merged into single bouts. Numerous properties of swim bouts were calculated. The maximum speed of a bout was determined from the largest displacement across two frames during the bout. Displacement across each pair of frames at speeds above 4 mm/sec was summed to find net bout displacement. Pitch angle distributions were computed pitch angles during IBIs. To calculate probability distributions of parameters, data from all bouts was pooled in each group (1240/1780 bouts and 3105/7392 IBI pitch measurements for mutant/control). Number of bins for plotting probability distributions satisfy Sturges’ rule (13 bins for speed and displacement; 15 bins for posture). Confidence intervals were estimated by resampling data 1000 times with a sample size 1/5 of that of the group (mutant or control) with fewer number of measurements (n = 248 for speed and displacement; n = 621 for IBI posture). To determine statistical significance, two-sample Kolmogorov-Smirnov tests were applied to original measurements (speed, displacement, and posture).

### Quantification and statistical analysis

#### Cell counting

Cell counting was performed using Fiji ^99^ or Imaris (Bitplane). Statistical analyses of cell counting were carried out using Prism 9 software with two-tailed Student’s t-tests unless otherwise noted. Differences were considered to be significant when p values were below 0.05. Graphs of quantitative data are plotted as means with standard error of mean (SEM) as error bars.

#### Muscle domain fluorescence

Domain fluorescence was measured using Fiji ^99^. Z-stacks of spinal muscle were cropped to one myotome and rotated 90°. A maximum intensity projection of the transverse view of the GFP (Prdm16+) channel was made. An ROI was drawn for each muscle quadrant, as defined in Figure 6A, per ^31^. The mean pixel intensity was calculated for each domain and normalized to domain IV.

#### Sparse label analysis

Hypothesis testing was performed by bootstrapping. First, a null distribution of cell types was estimated by resampling with replacement. p values were then computed explicitly as the fraction of times that the number of cells drawn from the null distribution met or exceeded the observed number. Each p value corresponds to n=10,000 such simulations. The null distributions were either the distributions of all motor neurons, derived from counts in ^31^ (26 types, n=303 cells), or the distribution of *prdm16+* motor neurons (n=51 cells).

## MOVIES

**Movie 1: examples of swimming after loss of *mecom***

The movie shows four brief series of swim bouts from control larvae raised without swim bladders. Next, four series of swim bouts from *mecom*-/- siblings are shown. The *mecom* -/- larvae can maintain a dorsal-up orientation.

**Movie 2: examples of swimming after loss of *prdm16***

The movie shows six series of swim bouts from *prdm16*-/- larvae. In contrast to *mecom*-/- larvae, prdm16 mutants cannot maintain a dorsal-up orientation. Instead, they adopt a wide range of postures in pitch (nose-up/nose-down) and roll axes, and “somersault” as in the last few series of bouts.

## ACKNOWLEDGMENTS

Research was supported by the National Institute on Deafness and Communication Disorders of the National Institutes of Health under award numbers R00DC012775, R56DC016316, R01DC017489 to DS, and F31DC019554 to KRH, and the National Institute of Neurological Disorders and Stroke under award numbers F31NS110235 to KPD, F99NS129179 to DG, T32NS086750 to KD and DG, and R35NS116858 to JSD, by the Leon Levy Foundation to YZ, by the Irma T. Hirschl/Monique Weill-Caulier Trust, by the Brain Research Foundation, and by the Dana Foundation to DS, and by Cancer Center Support Grant P30CA016087 to the NYU Langone Health Genome Technology Center. Transgenic support came from the NIG Scientist and the National BioResource Project from the Ministry of Education, Culture, Sports, Science and Technology of Japan. We thank members of the Knaut, Kucenas, McLean, and Liddelow labs for technical assistance with *in situ*, antibody staining, motor neuron classification, and especially Philip Hasel for single cell sequencing analysis. We thank members of the Dasen, Liddelow, Nagel, and Schoppik labs for support and advice.

## AUTHOR CONTRIBUTIONS

Conceptualization: KPD, Methodology: KPD, DS, JD Investigation: KPD, HH, DG, MK, YZ, and KH Writing: KPD, DS, JD Funding Acquisition: KPD, DS, JD, Supervision: DS, JD.

---

**Figure S1:**
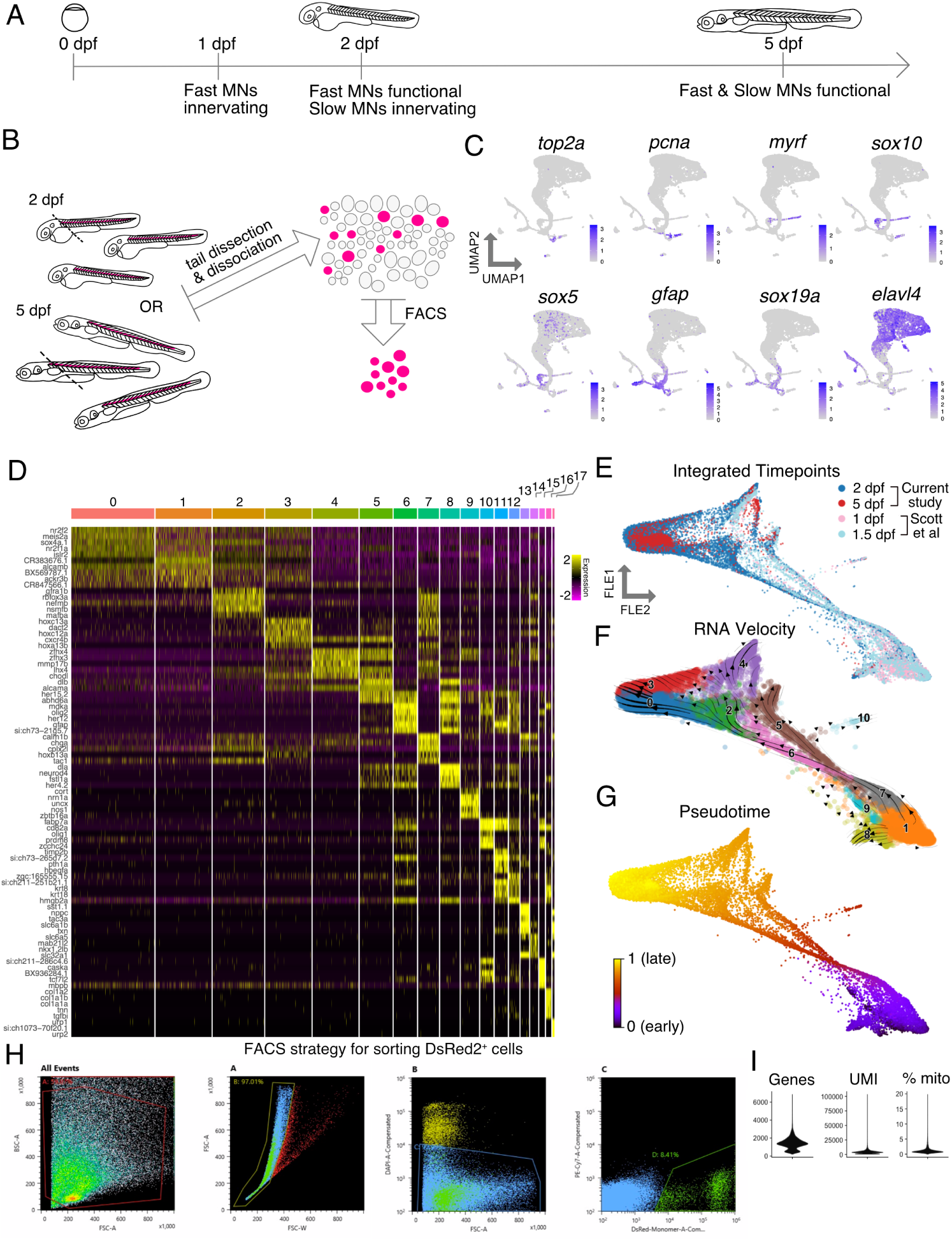
Using single cell RNA sequencing to evaluate motor neuron heterogeneity over time. (A) Timeline of zebrafish MN development indicating major timepoints of functional subtype maturation. (B) Schematic representing the scRNAseq experimental set up; dashed line represents dissection cut; magenta represents DsRed2^+^ cells from *Tg(olig2:DsRed2)* fish. FACS, fluorescence activated cell sorting. See FACS method in (H) (C) Feature plots of marker genes for major cell types-proliferating cells (*top2a*^+^, *pcna*^+^), oligodendrocytes (*myrf* ^+^, *sox10*^+^, *sox5*^+^), radial glia (*gfap*^+^, *sox19a*^+^, *pcna*^-^), interneurons (*elavl4*^+^, *olig2*^-^), and pMN progenitors (*pcna*^+^, *sox19a*^+^, *gfap*^-^) using UMAP coordinates in Figure 1B. Purple indicates expression. (D) Heatmap showing top five marker genes for each cluster; Each column is a cell; cells are grouped by cluster indicated by number and color; High expression shown in yellow. (E) Dimensionality reduction of merged 2 dpf and 5 dpf MN samples from this study with 1 dpf and 1.5 dpf MN samples from ^39^ shown via a force-directed graph (FLE). (F) FLE graph of merged samples with RNA velocity streamlines overlaid depicting flow of cell states over time based on splice ratios. (G) RNA velocity calculation of pseudotime with black as the youngest cells within the progenitor clusters and yellow indicating oldest clusters of MNs. (H) FACS gating strategy for sorting DsRed2^+^ cells. Gate A isolated cells within the size range to exclude debris. Gate B excluded large cells or doublets. Gate C excluded DAPI^+^ dead cells. Gate D isolated DsRed^+^ cells based on a negative control sample (not shown). (I) Violin plots of quality control metrics. Genes is the number of features per cell. UMI is the number of unique molecular indicator counts per cell. % mito is the percentage of mitochondrial counts counted for each cell.

**Figure S2:**
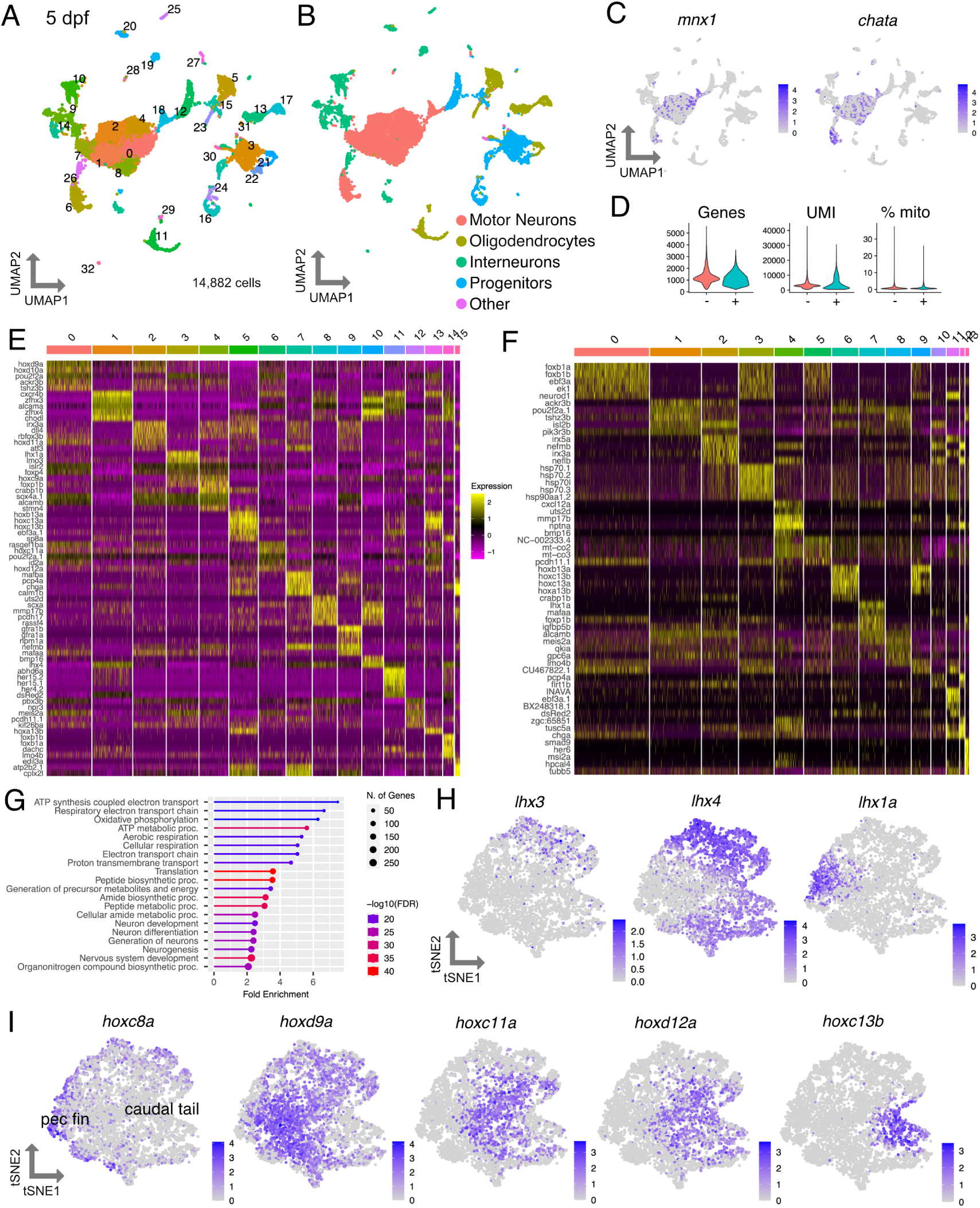
Molecular heterogeneity of motor neurons during development and at functional maturity. (A) UMAP representation of 14,882 *olig2*^+^ cells at 5 dpf from 580 larvae; Each dot represents one cell; 32 cell clusters are represented by coloring. (B) Major cell types represented in the UMAP. (C) Feature plots of marker genes for MNs. Purple indicates expression. (D) Violin plots of quality control metrics. Genes is the number of features per cell. UMI is the number of unique molecular indicator counts per cell. % mito is the percentage of mitochondrial counts counted for each cell. Note this experiment sorted for GFP and DsRed2 from *Tg(prdm16::GFP)*;*Tg(olig2:DsRed2)* fish as represented in Figure S7A. The two samples are represented by - (DsRed^+^ GFP^-^) and + (DsRed^+^ GFP^+^). The samples were combined for downstream analysis of the entire DsRed^+^ population. Asterisk indicates significantly differential gene expression. (E) Heatmap indicating top 5 marker genes for each cluster from 2 dpf MN tSNE in Figure 2A. High expression indicated in yellow. (F) Heatmap indicating top 5 marker genes for each cluster from 5 dpf MN tSNE in Figure 2B. (G) Gene Ontology analysis of all 2 dpf MN cluster markers; results from ShinyGO ^93^. (H) Feature plots of *lhx* gene family expression in 2 dpf mature MN tSNE. (I) Feature plots of *hox* gene family expression in 2 dpf mature MN tSNE.

**Figure S3:**
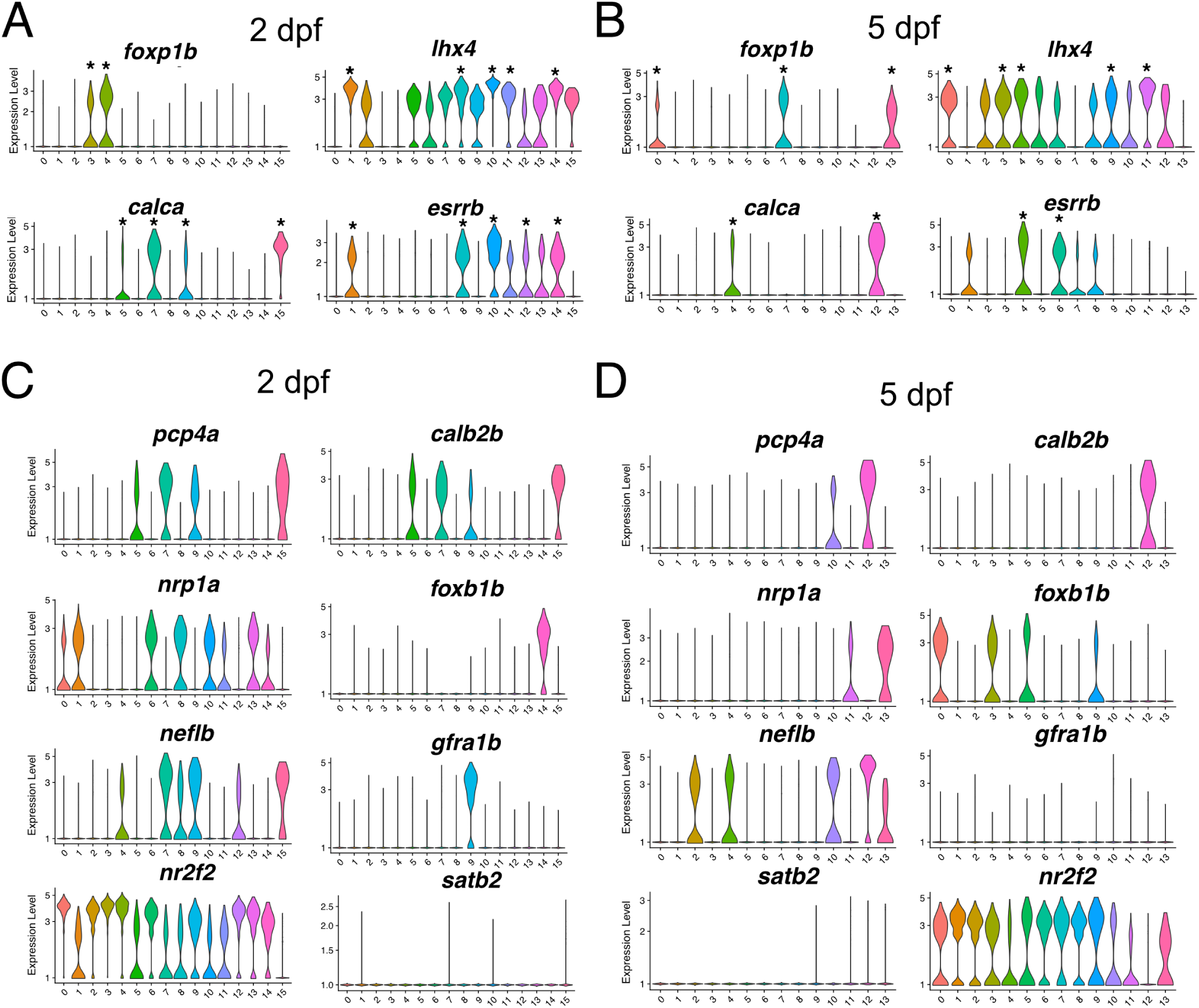
Distributions of expression level for select genes in MNs at 2 and 5 dpf. (A) Marker genes at 2dpf; clusters with significant differential expression marked with asterisks (B) Marker genes at 5dpf; clusters with significant differential expression marked with asterisks (C) Expression levels for select genes in each cluster at 2dpf (D) Expression levels for select genes in each cluster at 5dpf

**Figure S4:**
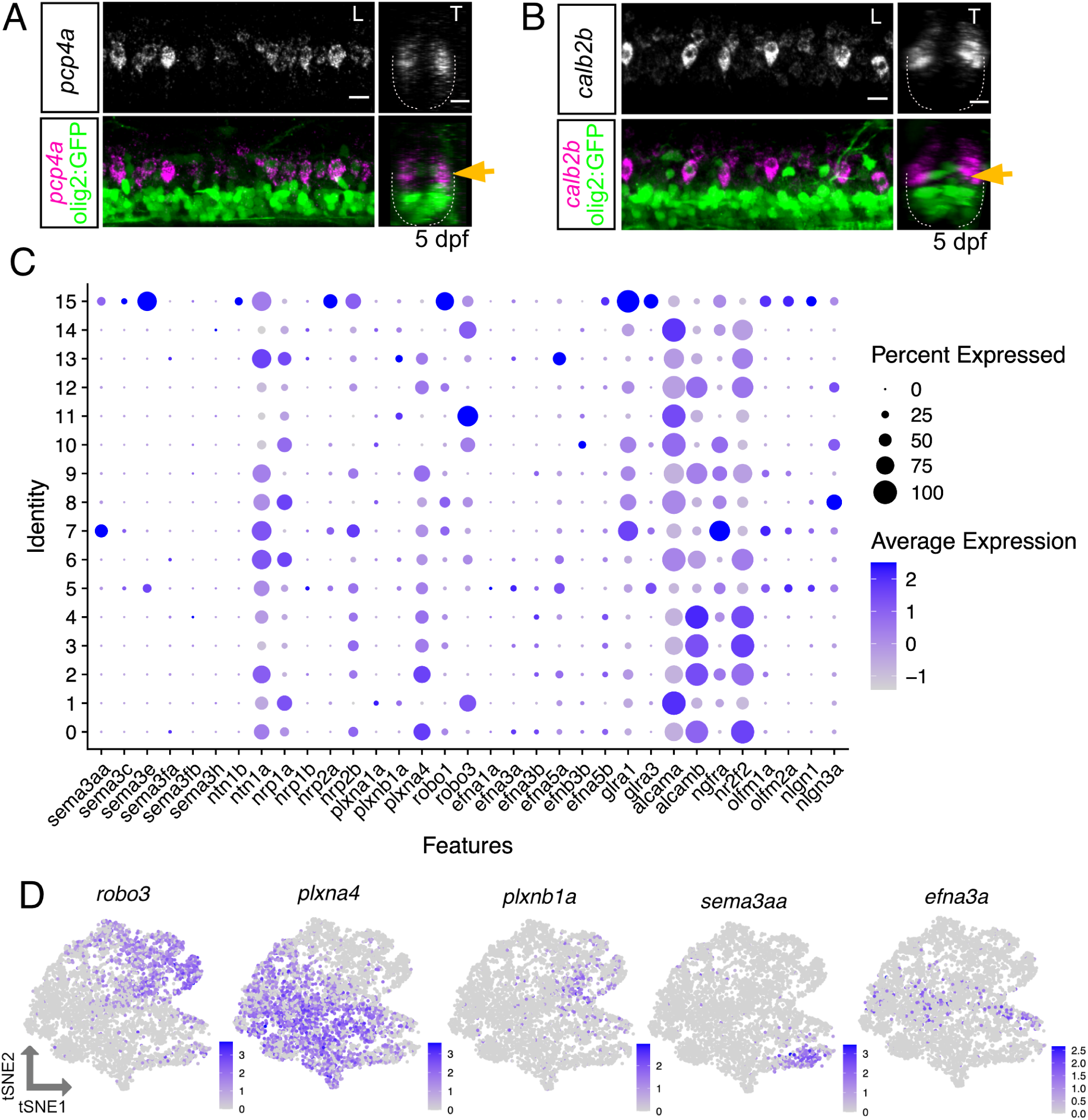
Evaluating developmental and sustained motor neuron subpopulation markers. (A-B) FISH for fast MN subpopulation markers at 5 dpf. Gold arrow indicates dorsoventral localization of FISH expression within MN population. Left image (L) is lateral view and right image (T) is transverse view. Scale bars, 10 µm. (C) Dot plot indicating expression of guidance cues within 2 dpf mature MN clusters in Figure 2A. (D) Feature plots showing example guidance cues from (C) with cluster-specific expression within the 2 dpf MN tSNE as in Figure 2A.

**Figure S5:**
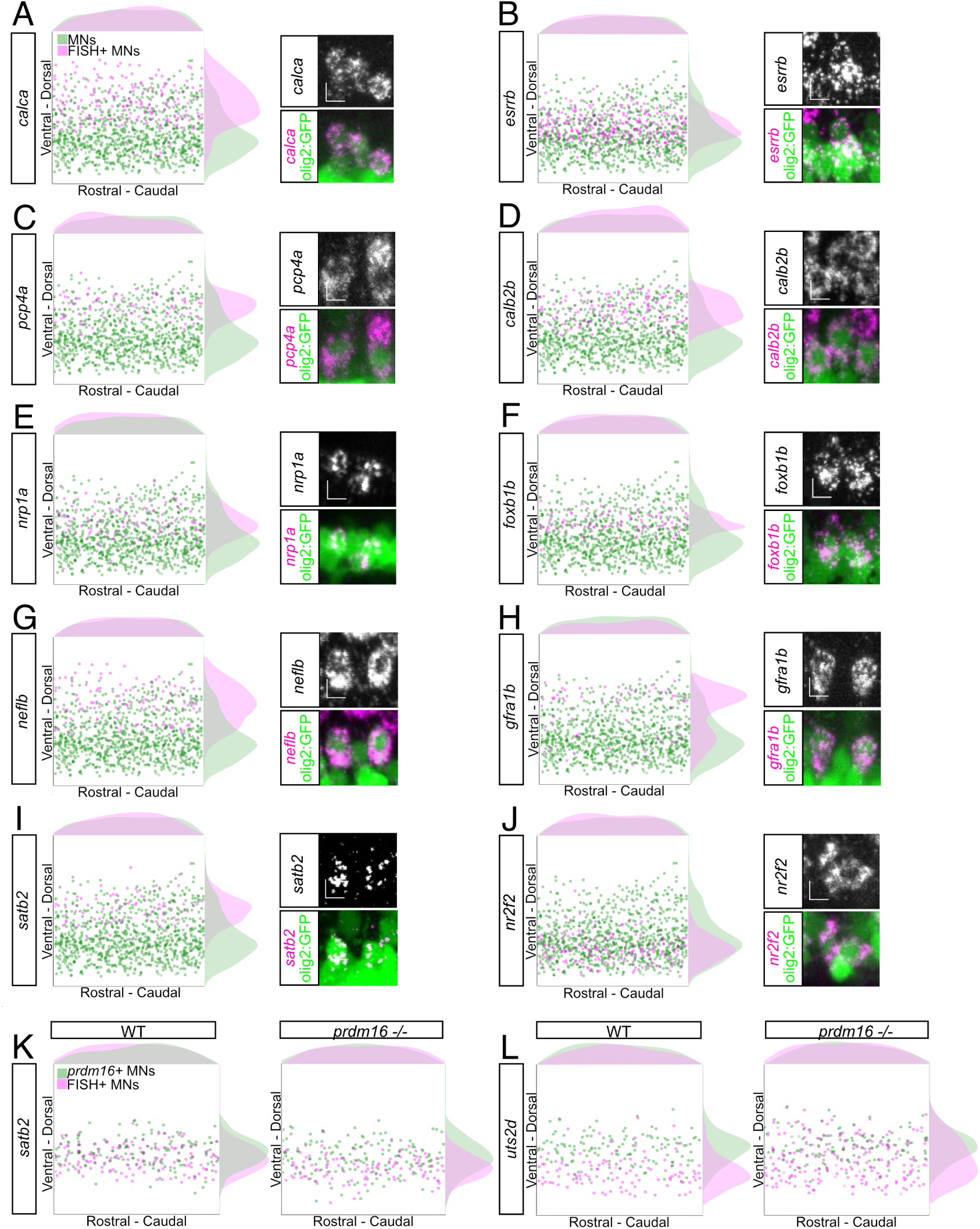
Further characterization and quantification of motor neuron functional subtype-restricted markers. (A-J) Lateral soma position plots for FISH probes, shown in Figures 3C to 3J, and all other MNs quantified from four larvae per probe at 2 dpf. B displays *esrrb* soma positions from seven larvae. Histograms on the top and sides of the plots indicate the distribution of cells across the axes. Green indicates GFP^+^;Olig2^+^ cells. Magenta indicates GFP^+^;FISH^+^ cells. To the right of each plot, a zoomed in image of FISH^+^;GFP^+^;Olig2^+^ somata highlights a MN expressing each specified probe. Scale bars, 5 µm. The percentage of total MNs for each gene is as follows: *calca*:37.3%, *esrrb*: 34.6%, *pcp4a*: 16%, *calb2b*: 26.4%, *nrp1a*: 27.5%, *foxb1b*: 25.8%, *neflb*: 22%, *gfra1b*: 18.6%, *satb2*: 13.3%, *nr2f2*: 43.6%. (K-L) Lateral soma position plots for FISH probes and *prdm16*^+^ MNs, shown in Figures 7C and 7E, quantified from four larvae per probe per condition at 5dpf. Green indicates GFP^+^;Prdm16^+^ cells. Magenta indicates GFP^+^; FISH^+^ cells.

**Figure S6:**
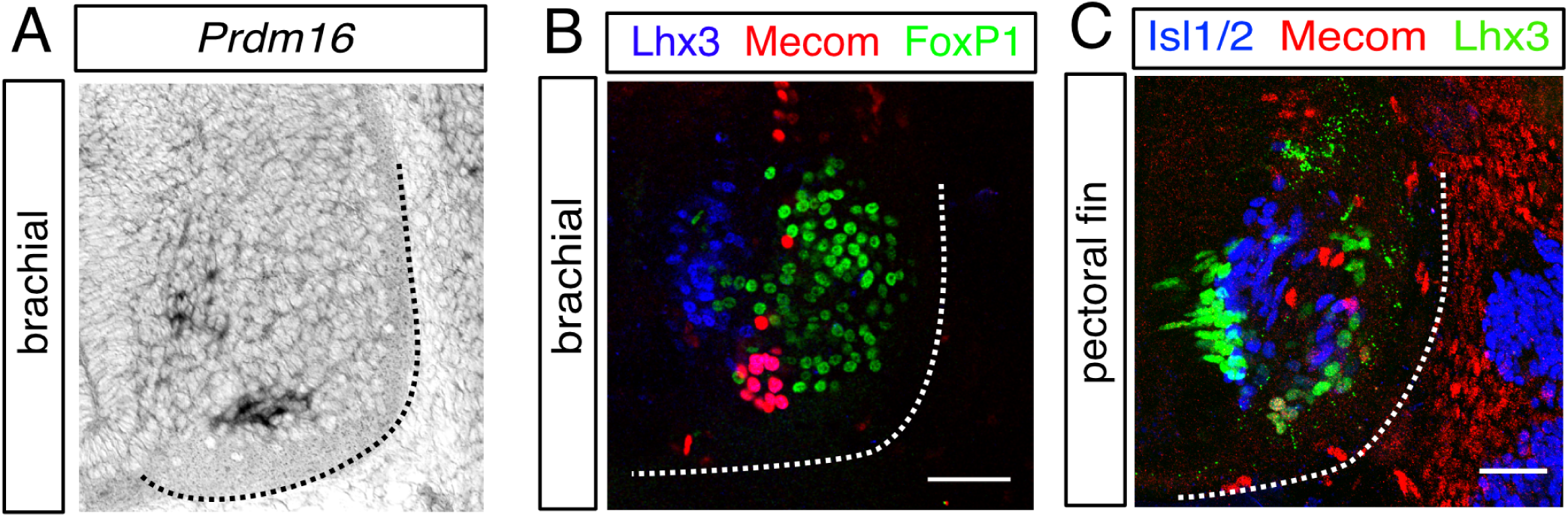
Conserved expression of *prdm16* and *mecom*. (A) *in situ* hybridization of *Prdm16* in the mouse ventral horn at E12.5. Black dotted line indicates ventral horn boundary. (B) Antibody stain of Evi1/Mecom expression within the ventral horn of chick spinal cord at HH27. White dotted line indicates ventral horn boundary. Scale bar, 50 µm. (C) Antibody stain of Evi1/Mecom expression within the ventral horn of skate spinal cord at stage 29. White dotted line indicates ventral horn boundary. Scale bar, 50 µm.

**Figure S7:**
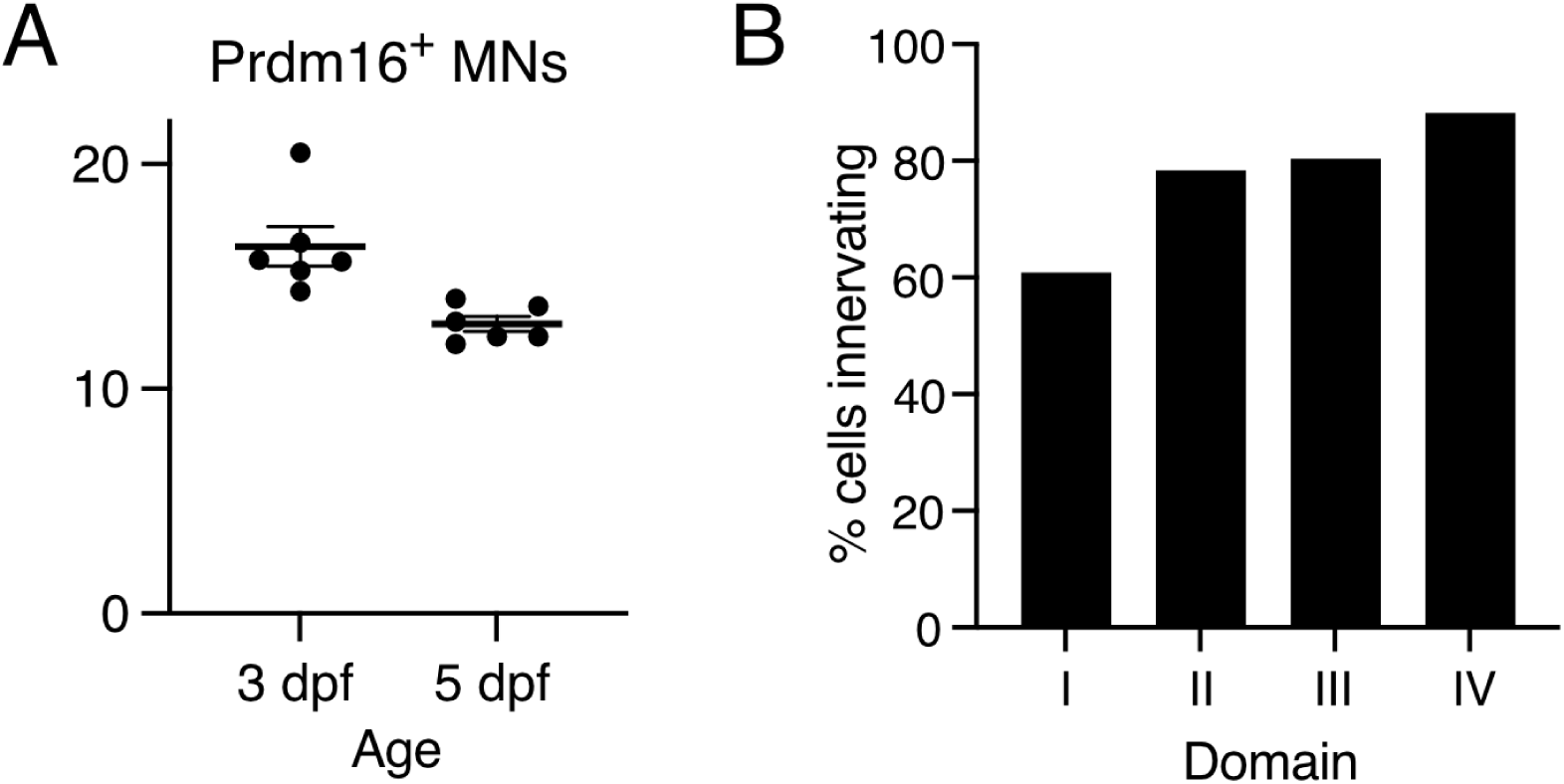
Characterizing Prdm16-expressing motor neuron population. (A) Quantification of GFP^+^ and DsRed2^+^ MNs in *Tg(prdm16::GFP)*;*Tg(olig2:DsRed2)* fish at 3 dpf and 5 dpf. Line indicates average with SEM. (B) Percentage of cells from sparse label experiment in Figure 5E that innervated each dorsoventral muscle quadrant.

**Figure S8:**
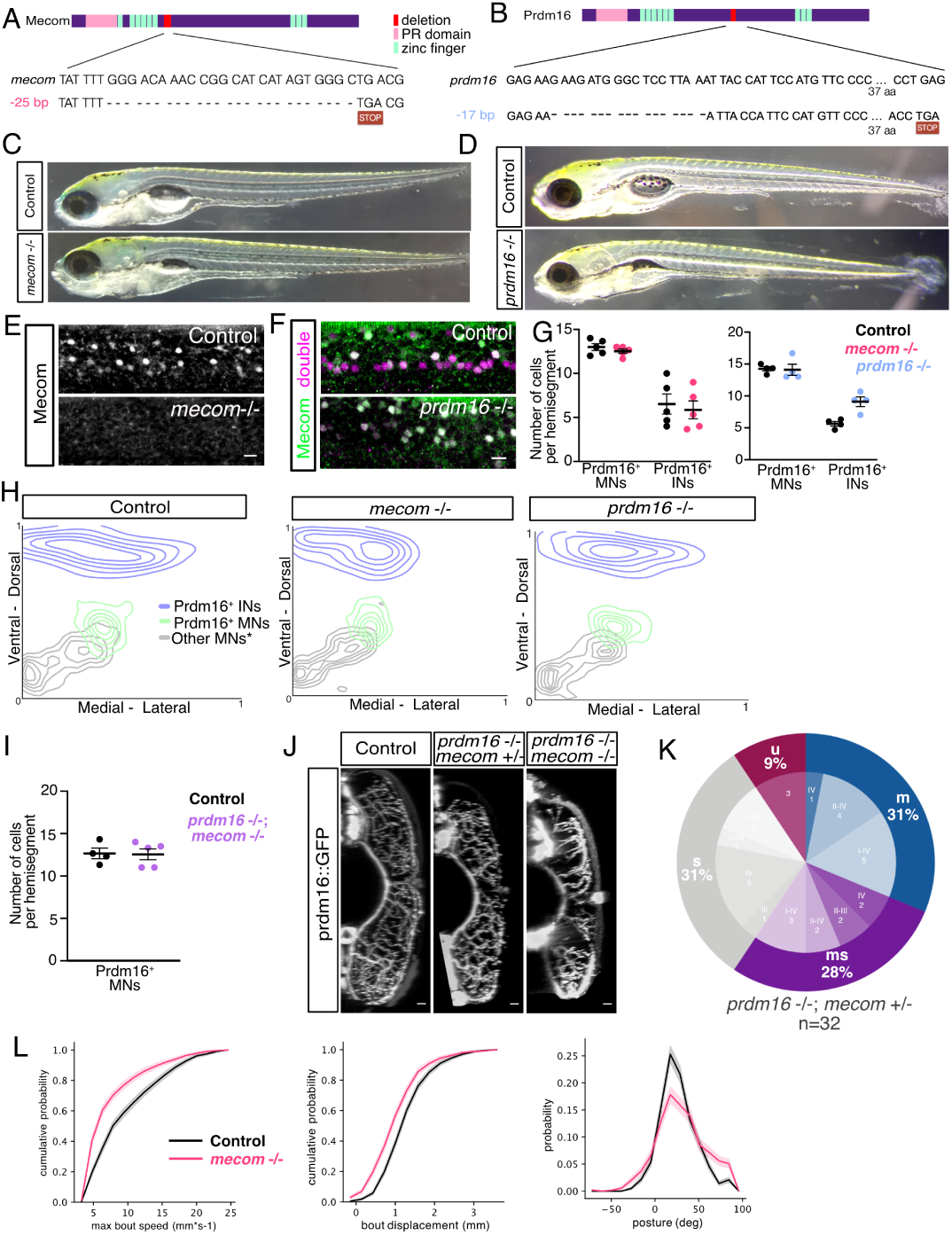
Assessment of *prdm16* and *mecom* mutants. (A) Schematic representing CRISPR mutation created in *mecom*. Pink represents the PR domain. Green represents the zinc fingers. Red represents the location of the mutation. Inset shows deleted nucleotides and resulting premature stop codon. (B) Schematic representing CRISPR mutation created in *prdm16*. Similar mutation (22 bp deletion at same location) was also created in the *Tg(prdm16:GAL4)* mutant line (not shown). (C) Images of *mecom* mutants and control siblings at 5 dpf. (D) Images of *prdm16* mutants and control siblings at 5 dpf. (E) Antibody staining for Mecom in 2 dpf wildtype embryos and *mecom* mutant larvae. Scale bar, 10 µm. (F) Antibody staining for Mecom and Prdm16/Mecom in 2 dpf in control and *prdm16* mutant larvae. Absence of Prdm16/Mecom^+^;Mecom^-^ cells in mutant indicate loss of Prdm16^+^ cells. Double in magenta indicates Prdm16/Mecom double antibody. *prdm16*-/-;*mecom*-/- double mutant larvae had complete loss of Prdm16/Mecom antibody staining demonstrating loss of both proteins (data not shown). Scale bar, 10 µm. (G) Quantification of Prdm16^+^ MNs and Prdm16^+^ interneurons (INs) from *Tg(prdm16::GFP)*;*Tg(olig2:DsRed2) mecom* mutant, *prdm16* mutant, and control larvae at 5 dpf. Line indicates average with SEM. (H) Contour map of the position of Prdm16^+^ INs (blue), Prdm16^+^ MNs (green), and Prdm16^-^ MNs (gray) from the transverse view of control (n=5), *mecom* mutant (n=5), and *prdm16* mutant (n = 3) *Tg(prdm16::GFP)*;*Tg(olig2:DsRed2)* larvae at 5 dpf. Highest density of cells represented by the smallest circle. While other MNs (gray) counts likely include some progenitors or oligodendrocytes, these were largely excluded based on morphology and localization. (I) Quantification of Prdm16*+* MNs from *Tg(prdm16d22::GFP)*;*Tg(olig2:DsRed2)* double mutant and control larvae at 5 dpf. Line indicates average with SEM. (J) Transverse view of the zebrafish tail showing Prdm16^+^ MN muscle innervation in *Tg(prdm16::GFP)* control, *prdm16*-/-; *mecom*+/- mutant, and *prdm16*-/-; *mecom*-/- double mutant larvae at 5 dpf. Scale bars, 10 µm., control is same as in Figure 6A (K) Pie chart showing subtype proportions labeled in*Tg(prdm16:GAL4)* sparse label for *prdm16*-/-; *mecom*+/- mutant larvae (L) Left plot: Cumulative probability of the maximum speed reached during each bout. The left shift of the mutant curve demonstrates the increased likelihood *mecom* mutants will swim at slower speeds than control larvae. (1240/1786 bouts for 96 control/120 mutant larvae; K-S p=5.14e-44) Middle Plot: Cumulative probability of the distance swam during each bout. Left shift of the mutant curve shows *mecom* mutants are more likely to swim shorter distances. (1240/1786 bouts for control/mutant; K-S p=2.79e-26) Right Plot: The probability distribution of postures assumed by larvae during the inter-bout interval (when fish is not swimming). *mecom* mutants are more likely to swim at exaggerated nose up postures. (3105/7392 sampled pitch timepoints from 594/1070 inter-bout interval for control/mutant; K-S p=1.88e-22); All data bootstrapped, line = mean, shaded area = SD.

**Figure S9:**
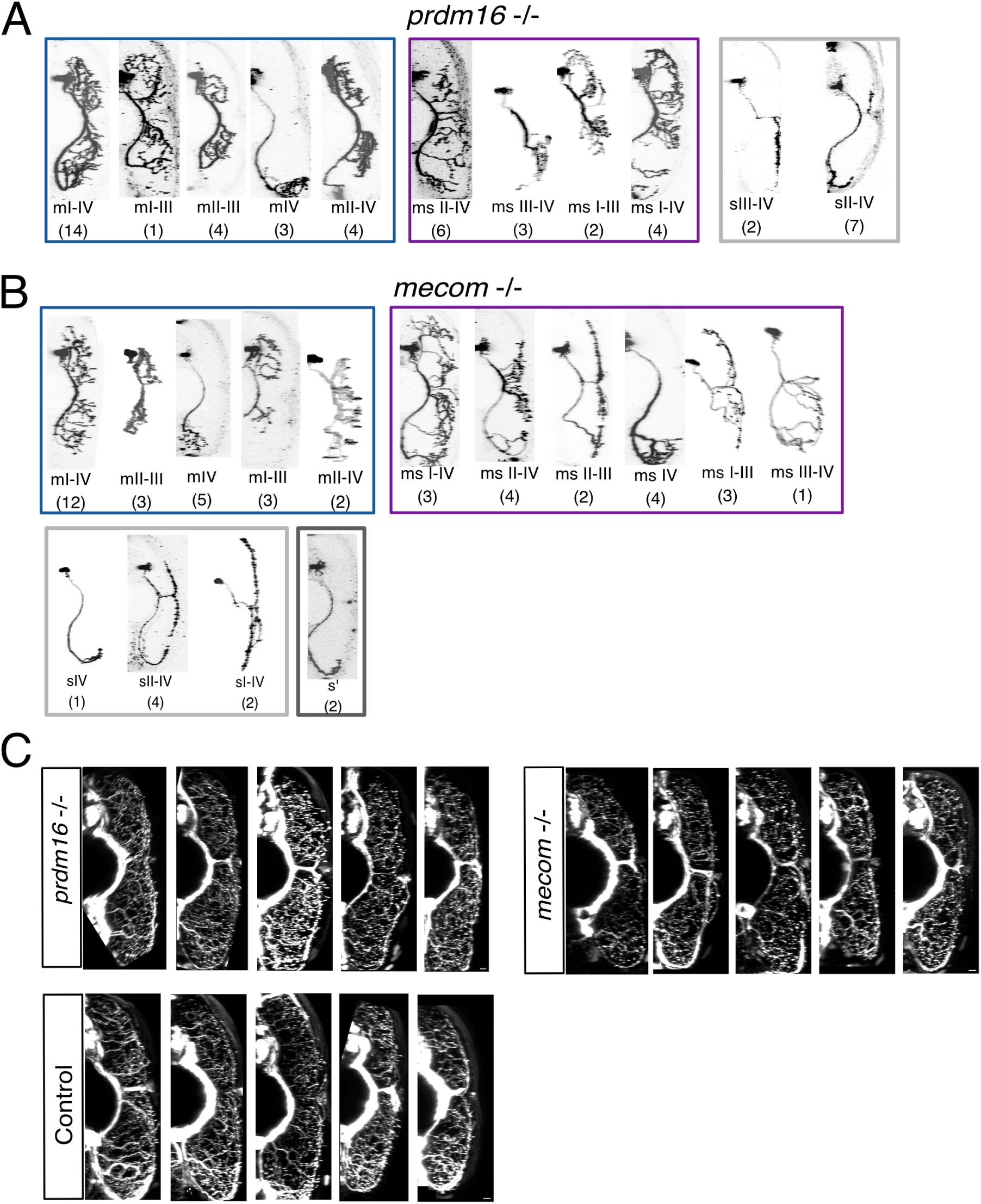
Changes in axonal innervation in *prdm16* and *mecom* mutants. (A) Example images of all sparsely labeled subtypes in *prdm16* mutant data set (cf. Figure 6F). (B) Example images of all sparsely labeled subtypes in *mecom* mutant data set (cf. Figure 6E). Number in parentheses indicates number of times subtype was labeled. (C) Additional examples of transverse view of the zebrafish tail showing Prdm16+ MN muscle innervation in *Tg(prdm16::GFP)* control, *mecom* mutant, and *prdm16* mutant larvae at 5 dpf. Scale bars, 10 µm.

**Figure S10:**
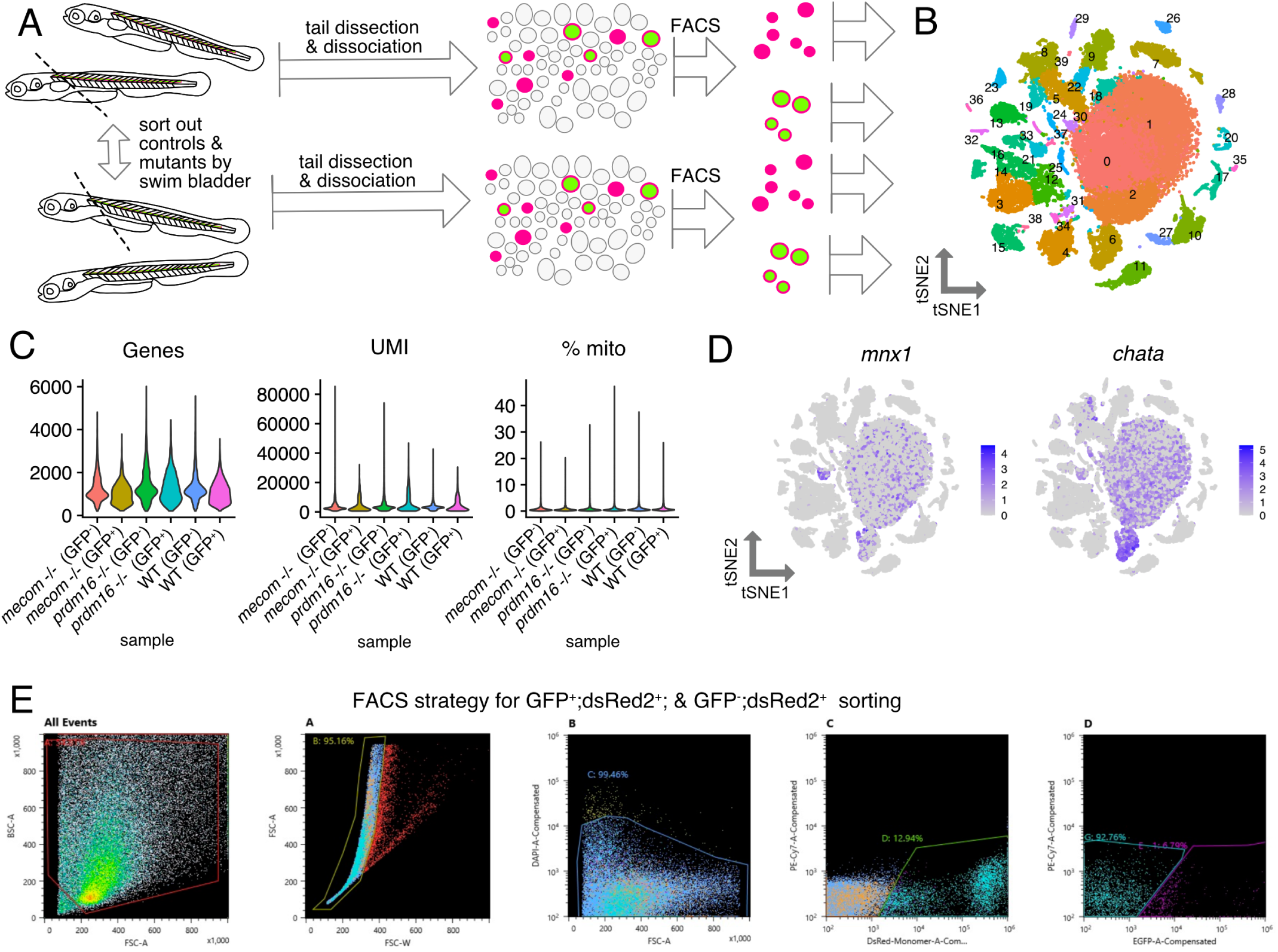
Single cell RNA sequencing of *prdm16* and *mecom* mutant motor neurons. (A) Schematic of scRNAseq experiment with *Tg(prdm16::GFP)*;*Tg(olig2:DsRed2)* WT, *mecom* mutant, and *prdm16* mutant larvae. (B) tSNE map of all six MN samples combined for analysis: WT GFP^-^;DsRed2^+^ cells and WT GFP^+^;DsRed2^+^ cells from 560 larvae, *mecom* mutant GFP^-^;DsRed2^+^ cells and *mecom* mutant GFP^+^;DsRed2^+^ cells from 170 larvae, *prdm16* mutant GFP^-^;DsRed2^+^ cells and *prdm16* mutant GFP^+^;DsRed2^+^ cells from 180 larvae. (C) Violin plots of quality control metrics demonstrating the quality and depth of sequencing is comparable between samples. Genes is the number of features per cell. UMI is the number of unique molecular indicator counts per cell. % mito is the percentage of mitochondrial counts counted for each cell. (D) Feature plots for MN markers for tSNE in B. (E) FACS gating strategy for sorting GFP^+^;DsRed2^+^ and GFP^-^;DsRed2^+^ cells. Gate A isolated cells within the size range to exclude debris. Gate B excluded large cells or doublets. Gate C excluded DAPI^+^ dead cells. Gate D isolated DsRed^+^ cells based on a negative control sample (not shown). Gate E isolated GFP^+^ cells and Gate G isolated GFP^-^ cells based on a negative control sample (not shown).

